# Conformational gating at histidine junctions coordinates proton translocation in respiratory complex I

**DOI:** 10.64898/2026.08.12.744383

**Authors:** William Fisher, John J. Wright, Man Nok Choy, Guilherme M. Arantes, Robert A. Waddell, Daniel N. Grba, Judy Hirst

**Author notes:** These authors contributed equally (William Fisher, John J. Wright). Authors for correspondence (JJW), (JH).

## Abstract

Respiratory complex I, a central enzyme in cellular metabolism, converts the free energy of NADH oxidation into a transmembrane proton-motive force to drive ATP synthesis, but the molecular mechanisms by which it couples redox catalysis to vectorial proton translocation remain unresolved. Here, we present high-resolution cryo-EM structures of complex I from *Bos taurus* captured under conditions designed to change the protonation states of residues in the membrane domain. Our structures reveal conformational rearrangements at key pathway junctions that reconfigure proton-transfer connections. In ND5, helical rearrangements switch the connectivity of histidine-248 between proton-uptake and proton-output pathways. In ND4, rotameric changes of histidine-220 alternately enable proton uptake or lateral proton transfer along the membrane domain. Combined with molecular simulations, our structures define gating mechanisms that impose directionality on proton transfer reactions and provide a framework for proton-coupled energy transduction in complex I.

## Main text

Respiratory complex I (NADH:ubiquinone oxidoreductase) is central to mitochondrial metabolism^1–4^. It captures the free energy from ubiquinone reduction by NADH to drive protons across the inner mitochondrial membrane, contributing to the proton-motive force (Δp) that drives ATP synthesis. It is also a locus of reactive oxygen species production, a key regulator of mitochondrial redox balance, and its dysfunctions are a leading cause of primary mitochondrial and associated neuromuscular disorders^5,6^. In recent years, advances in cryo-electron microscopy (cryo-EM)^7^ have transformed our knowledge of complex I through high-resolution structures solved in multiple states from mammalian species^8–11^ as well as diverse model organisms^12–17^.

Mammalian complex I contains fourteen highly conserved catalytic ‘core’ subunits and 31 ‘supernumerary’ subunits with roles in assembly, stability, and regulation^18,19^. Together they form the redox-active hydrophilic domain and the proton-pumping membrane domain of the ∼1 MDa L-shaped assembly. In the hydrophilic domain, electrons from NADH oxidation by a flavin mononucleotide are transferred via a chain of iron-sulphur (FeS) clusters to ubiquinone-10 (Q10), bound in a deep channel (the Q-site) at the domain interface^20–22^. Two hydrated networks of residues connect the Q-site to three distal antiporter-like subunits (ND2, ND4, ND5) in the membrane domain, where proton pumping occurs^8,11,15,23,24^. The Glu-dominated ‘E-channel’ connects the Q-site to the ‘central axis’, which crosses each antiporter-like subunit via a Glu–Lys ion pair (between transmembrane helix (TMH) 5 and TMH7), a subunit-dependent series of intervening ionizable residues (notably Lys and His), and a ‘terminal’ Glu/Lys residue on TMH12^25–27^. Proton-uptake pathways from the matrix have been proposed to meet the central axis mid-way across each subunit, at TMH8-H248 in ND5, TMH7b-H220 or TMH8-K237 in ND4, and TMH7b-T119 or TMH8-K135 in ND2^8,12,15^, whereas only a single proton-output pathway to the intermembrane space (IMS) has been identified, from the terminal Lys in ND5^11,15^ (Fig. 1a).

**Fig. 1.**
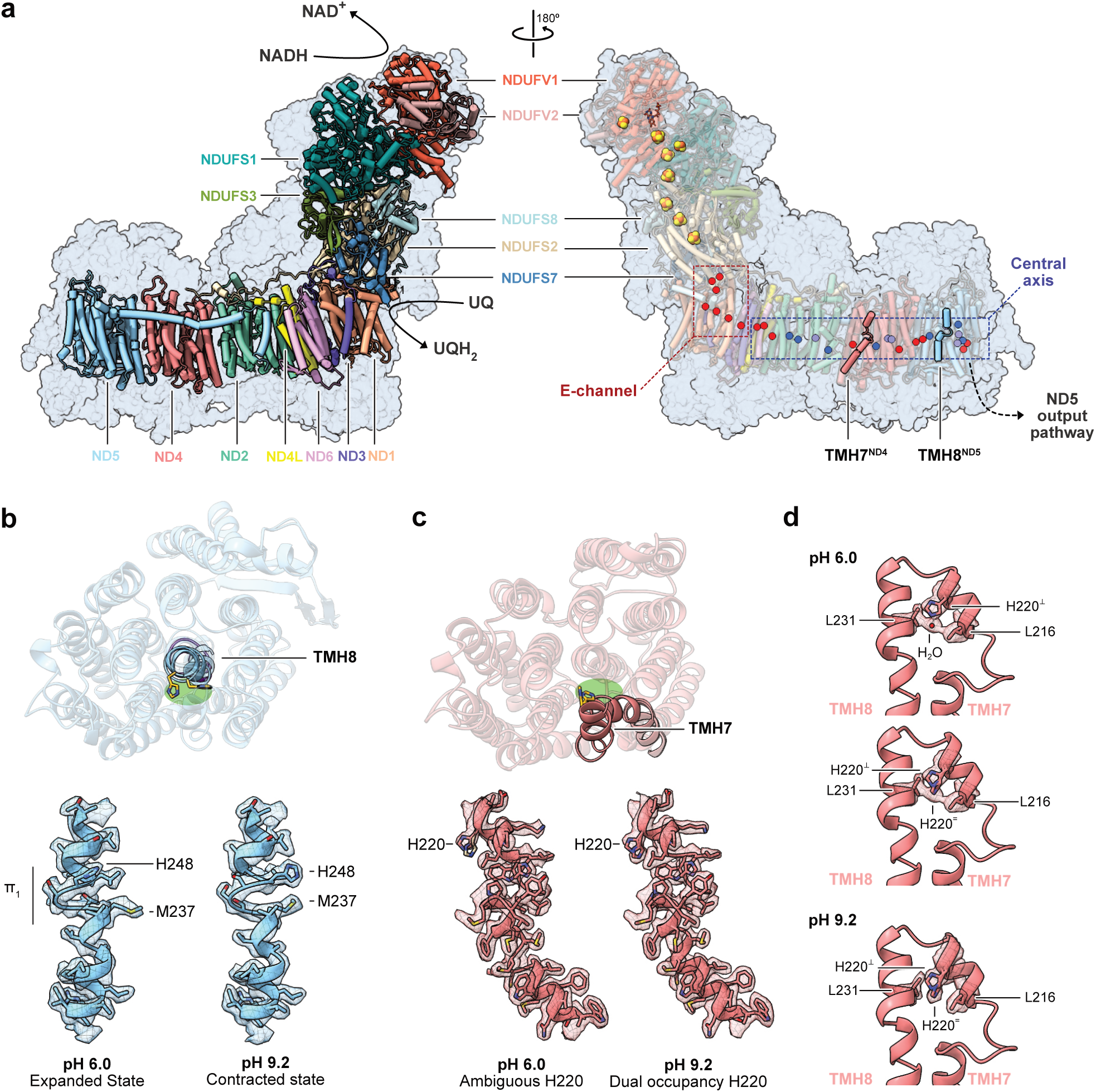
Complex I structures at pH 6.0 and G.2 reveal conformational changes in subunits ND4 and ND5. (**a**) Architecture of *B. taurus* complex I at pH 6.0. Core subunits are shown as tubes with the pale blue surface of the supernumerary subunits. The catalytic cofactors (flavin mononucleotide and FeS clusters) are shown on the right with Glu (red), Lys (blue) and His (pale purple) residues shown as spheres along the central axis. Helices with variations between structures at pH 6.0 and 9.2 are indicated. The structure is in the closed/A state. (**b**) The position and conformation of TMH8^ND5^ in the pH 6.0 and 9.2 structures. Top, TMH8^ND5^ is shown top-down for pH 9.2, (contracted) and pH 6.0 (expanded). The positions of H248 are indicated, and the region it occupies between TMH8 and TMH7 is in green. Bottom, density maps and models for TMH8 in the expanded and contracted states. (c) The position and conformation of TMH7^ND4^ in the pH 6.0 and 9.0 structures. Top, the equivalent position between TMH8 and TMH7 is in green with the two conformations of H220 shown. Bottom, density maps and models for TMH7 at pH 6.0 and 9.2. (**d**) Analysis of the densities at the TMH7-8 turn in ND4 at pH 6.0 and 9.2. Densities are shown for H220, L216 and L237 with two alternative models for the ambiguous density of H220 at pH 6.0 (top, the perpendicular position with a water molecule and bottom, the dual conformation).

Although residues across the central axis are known to be essential for function, conformational variability within it is currently limited to H248^ND5^, where alternative conformations have been observed in structures from different species^12^. Subsequently, structural and computational work on the Multiple Resistance and pH Adaptation (MRP)-type H^+^/Na^+^ antiporter, an ancestral relative of complex I, suggested the homologous His switches between positions to enable and coordinate proton-pumping events^28–30^. Otherwise, static structures of complex I have provided little information on proton-transfer activity and dynamics within the networks, while computational approaches have indicated extensive internal hydration of the membrane domain^31^ and advocated for uptake and output channels in all three subunits^24,32,33^. The mechanisms coupling ubiquinone reduction to proton pumping, which must be underpinned by the routes of proton uptake and output, and the role of lateral proton transfer along the central axis, thus remain unknown^15,34–36^.

Here, we investigate how proton-transfer pathways are gated and controlled in mammalian complex I. The energetics and connectivity of these hydrated networks of ionizable residues are likely sensitive to their protonation states, which may be modulated by varying the solvent pH. However, most cryo-EM structures of complex I have been determined under conditions optimised for activity or stability, with a single previous attempt^12^ to probe pH-dependent effects limited by insufficient resolution and complications from regulatory transitions^37–41^. Here, we determine high-resolution structures (up to 2.2 Å) of a catalytically relevant state of complex I under defined pH conditions to investigate protonation-dependent conformations and hydrogen-bonding networks. The structures reveal conformational changes at central histidine residues in subunits ND4 and ND5 that switch their proton-transfer connections between proton-uptake pathways from the matrix and the ‘upstream’ (redox side) and ‘downstream’ branches of the central axis. We thereby begin to define how complex I gates proton-pumping events by using junction control points to alternately connect and segment proton-transfer pathways during catalysis.

## Results

### Complex I structures under defined pH conditions

To investigate how protonation-state changes influence the structure of mammalian complex I, we carried out cryo-EM analyses of complex I from bovine (*Bos taurus*) heart mitochondria at pH 6.0 and 9.2, to match existing structures at pH 7.5. The pHs were chosen using biochemical analyses to establish the pH range (6 –10) at which complex I remains stable and catalytically competent, alongside its typical bell-shaped pH-dependent activity profile (Extended Data Fig. 1).

Samples for cryo-EM were purified in *n*-dodecyl β-D-maltoside (DDM) using our standard protocol^42,43^, then pH-adjusted during size-exclusion chromatography (Extended Data Fig. 2a). Cryo-EM imaging yielded 803k (pH 6.0) and 722k (pH 9.2) complex I particles that were classified into three distinct states matching the closed/active (closed/A), open/deactive (open/D), and slack states described previously for bovine complex I^9,22,41^ with global resolutions of 2.2–2.5 Å (Supplementary Figs. 1–6, Supplementary Tables 1–2). The slack state, an inactive artifact of preparations in DDM^41,44^, accounted for 53% and 29% of particles at pH 6.0 and 9.2, compared with 20% at pH 7.5^9^, consistent with the cryo-EM sample activities (Extended Data Fig. 2b-c) and indications from 30 °C incubations of lower stability at pH 6.0 (Extended Data Fig. 2d). As the slack state increases at the expense of the open/D state (Extended Data Fig. 2c) our data suggest it is formed by pH (and DDM) induced instability of the open/D state. In contrast, the ‘turnover-ready’ closed/A and ‘dormant’ open/D states are physiologically relevant resting states, which comprise an ischemia-induced regulatory switch in the mammalian enzyme^37–41^. In the widely accepted model for the ‘deactive transition’ (Extended Data Fig. 3a), deactivation occurs slowly in the absence of substrates (ischemia) and activation occurs rapidly when substrates return and reinitiate catalysis (reperfusion). Although both rates depend on pH^45,46^, lack of substrates and low temperature should preclude interconversion during sample preparation, and indeed the proportion of closed/A particles is constant at 30–35% in all cases (Extended Data Fig. 2c). Conversely, medium-resolution cryo-EM analyses of complex I from *Ovis aries*^12^ reported varying proportions of closed/A particles (25 and 54% at pH 5.5 and 9.0) and a reversible, pH-dependent equilibrium between closed/A and open/D proposed, even at low temperatures and without substrates (Extended Data Fig. 3b). Suspecting the shift was caused by unknown differential stability effects, we used biochemical analyses to resolve the discrepancy (Extended Data Fig. 3c). Our data confirm the distribution is stable and unaffected by pH at 4 °C, supporting the extant model (Extended Data Fig. 3a).

Global comparisons of the closed/A and open/D structures revealed no large-scale pH-dependent changes (Extended Data Fig. 4a-b). Two local changes identified in subunits ND4 and ND5 in the closed/A structures (Extended Data Fig. 4c), also observed in the open/D structures, are the focus of this work. An additional change in the glutamate-rich TMH5-6^ND1^ loop in specifically the open/D structures is probably affected by the DDM bound nearby in the Q-site and not physiologically relevant (Extended Data Fig. 4d).

### Histidine-linked rearrangements in subunits ND5 and ND4

The differences identified in ND4 and ND5 are at two key histidines, TMH8-H248^ND5^ and TMH7-H220^ND4^ (Fig. 1). Their imidazole rings occupy equivalent physical spaces, on the central axis at the base of the proposed proton-uptake channels^8,12^ (Fig. 1b-c), although they are not conserved in sequence. No corresponding difference was identified in subunit ND2, where the matching position is occupied by a threonine in mammalian complex I.

The most pronounced rearrangement occurs at TMH8-H248^ND5^. At pH 9.2, residues 247-252 adopt a π-bulge configuration that we term ‘contracted’ whereas, at pH 6.0, they reorganize to a strikingly different configuration, termed ‘expanded’ to reflect the additional residue accommodated by the helical turn (Fig. 1b). The same change is observed in the open/D structures (Extended Data Fig. 5a). All previously reported mammalian and plant complex I structures adopt the contracted conformation, with the surprising exception of biguanide-bound bovine complex I^47^, while the expanded state is prevalent in bacterial, fungal and protist homologues (Supplementary Table 3). Structures at pH 6.5 consistently adopt the expanded conformation, whereas structures at higher pH occupy either state, implicating protonation events, as well as species specificity, in controlling the state occupancy. By changing the conditions and switching the conformation in structures from a single species, we demonstrate the conformations are dynamically interconvertible, supporting proposals from computational studies that H248^ND5^ moves to direct proton transfers along different pathways during catalysis^12,28,30^.

The structural variation observed at TMH7-H220^ND4^ comprises a more subtle sidechain reorientation, without changes to the helical framework (Fig. 1c). H220^ND4^ has been implicated in proton uptake^11,31^ and mutation of the equivalent residue in *Paracoccus denitrificans* abolishes catalysis^27^. At pH 9.2, the density for the H220^ND4^ sidechain clearly shows two conformations, termed ‘planar’ and ‘perpendicular’ for their orientation to the central axis (Fig. 1c-d). Although previously modelled in a single state, both conformations are represented in high-resolution (≤ 2.5 Å) structures, albeit often with ambiguous density (Supplementary Table 3). At pH 6.0, the density is less readily interpreted and consistent with either a mixture of conformations or a predominantly perpendicular rotamer, perhaps stabilized by a coordinating water at H220-Nε (Fig. 1d). Consistent with the latter, the planar conformation is not supported by the open/D structure (Extended Data Fig. 5b). Thus, while less clearly correlated to pH than in ND5, the pose of H220^ND4^ is also sensitive to the protonation environment.

### Structural basis of the ND5 conformational gate

The transition of TMH5^ND5^ between the contracted (pH 9.2) and expanded (pH 6.0) states is defined by a dramatic remodelling of the helical framework of residues 247-252 while the matrix-facing (241-246) and IMS-facing (253-262) segments remain fixed (Fig. 2a). From the contracted to expanded conformation: L247 advances a helical position and changes rotamer; H248 advances similarly, reorienting its sidechain from towards TMH5 to towards TMH10; S249 moves into the position vacated by S250, replacing its hydrogen-bonding interaction with an adjacent water. Residues S250, T251 and M252 then compress, packing three residues into the helical turn that previously only accommodated two. The transition fundamentally rewires the proton-transfer connectivity of the ND5 subunit.

**Fig. 2.**
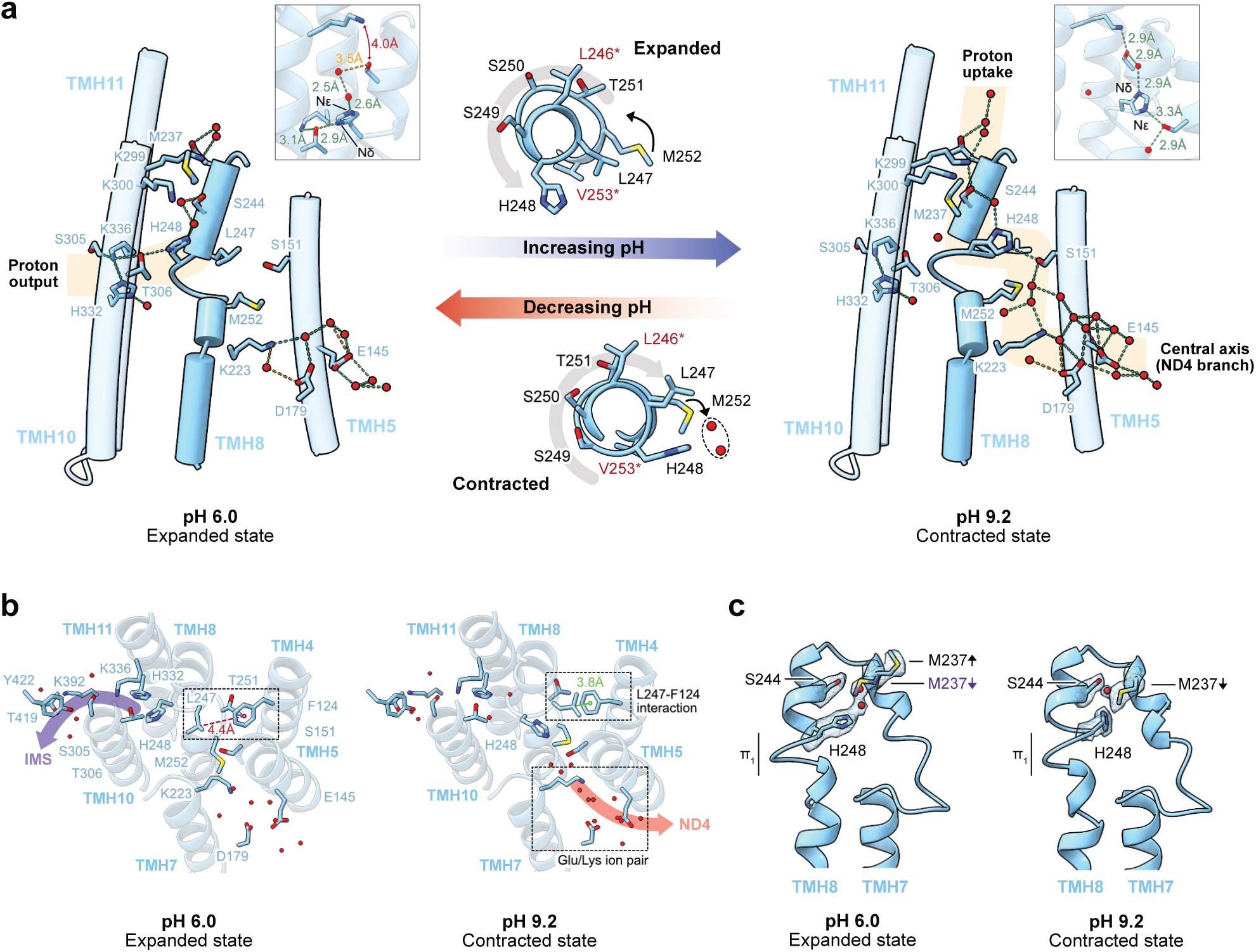
Protonation-dependent conformational changes in ND5 coordinate proton translocation. (**a**) pH dependent conformational changes around TMH8^ND5^ confer connectivity to the central axis, proton uptake and proton output channels. Arrows indicate the movement of residues in response to increasing and decreasing pH. Residues at the top (L246) and bottom (V253) of the expanding loop that do not shift position are labelled in red. Routes of connectivity are highlighted in pale orange. Insets show details of hydrogen bonding interactions made by H248. Hydrogen bonding interactions are shown as green (<3.4 Å) and orange (3.4 – 3.7 Å) dashes and ordered water molecules as red spheres. (**b**) Connectivity to the ND4-branch of the central axis is broken in the expanded state: repositioning of H248 severs the connection to the Glu/Lys ion pair and establishes connectivity with the proton-output pathway to the IMS. The hydrophobic interaction between TMH8-L247 and TMH4-F124 is disrupted upon helical expansion. (**c**) Expansion of TMH8 repositions TMH7-M237, which shifts from a downward conformation in the contracted state to an upward conformation in the expanded state. A region of continuous density between H248 and M237 is modelled as water molecules in the pH 6.0 structure (red spheres) but could alternatively be modelled as a dual conformation of upward- and downward-facing M237.

In the contracted conformation, H248 forms a junction between the proposed matrix-uptake pathway and the ‘upstream’ ND4-branch of the central axis. H248-Nε forms a hydrogen bond to TMH5-S151, which connects via a robust water network to the conserved Glu-Lys ion pair (E145-K223, augmented by D179) where the central axis enters from ND4 (Fig. 2a). H248-Nδ hydrogen bonds to a water, which bridges it to the proposed uptake-pathway residues TMH8-S244 and TMH10-K299. In this conformation, H248 is isolated from the ‘downstream’ output branch of the central axis, ∼10 Å away from TMH10-T306 with no continuous solvation to support proton transfer.

The transition to the expanded conformation severs the uptake and upstream connections and establishes a new connection to the proton-output pathway to the IMS, forming a hydrogen bond from the repositioned H248-Nδ to TMH10-T306, which in turn interacts with TMH11-K336 (Fig. 2a). Crucially, the connectivity switch is reinforced by formation of a hydrophobic barrier between H248 and the ND4-branch of the central axis, with L247 breaking its interaction with TMH4-F124, and M252 repositioning to displace the waters that previously linked S151 to the E145–K223 ion pair (Fig. 2b). The functional significance of this coordinated change is underscored by the association of F124 mutations with Leigh syndrome^48^. No clear connection from H248 towards the matrix is observed in the expanded conformation, though the local density requires careful interpretation. While two water molecules can be modelled between H248-Nε and TMH8-S244, the connection is obstructed by a rotamer shift in K299. The shift is induced by the TMH7-M237 sidechain adopting an “upward” pose, which is clearly absent in the contracted state (Fig. 2c). While the density in the expanded state is consistent with the upward pose, an alternative interpretation involving dual-occupancy (as noted in *P. denitrificans*^13^) would see a “downward” M237 occupying the density assigned to the two waters. In either scenario, M237 severs the link between H248 and the matrix-uptake pathway. Collectively, the coordinated transitions between the contracted and expanded states define H248 as a molecular gate that defines the proton-transfer connectivity at the junction of the central axis and matrix-uptake pathways, ensuring the directionality of proton translocation.

### The H220 rotameric gate in ND4

H220^ND4^ is located on TMH7, with its sidechain pointing towards TMH8, occupying the same region between TMH5 and TMH10 as H248^ND5^ (Fig. 1b-c). Our structures show the H220 imidazole ring toggles between planar and perpendicular conformations, defined by rotations around the Cα–Cβ and Cβ–Cγ bonds (Fig. 3a-b). By comparing our maps with previously determined high-resolution structures (Fig. 3c-d), we define the two rotamers as discrete functional states that re-route the proton-transfer connectivity of the ND4 subunit.

**Fig. 3.**
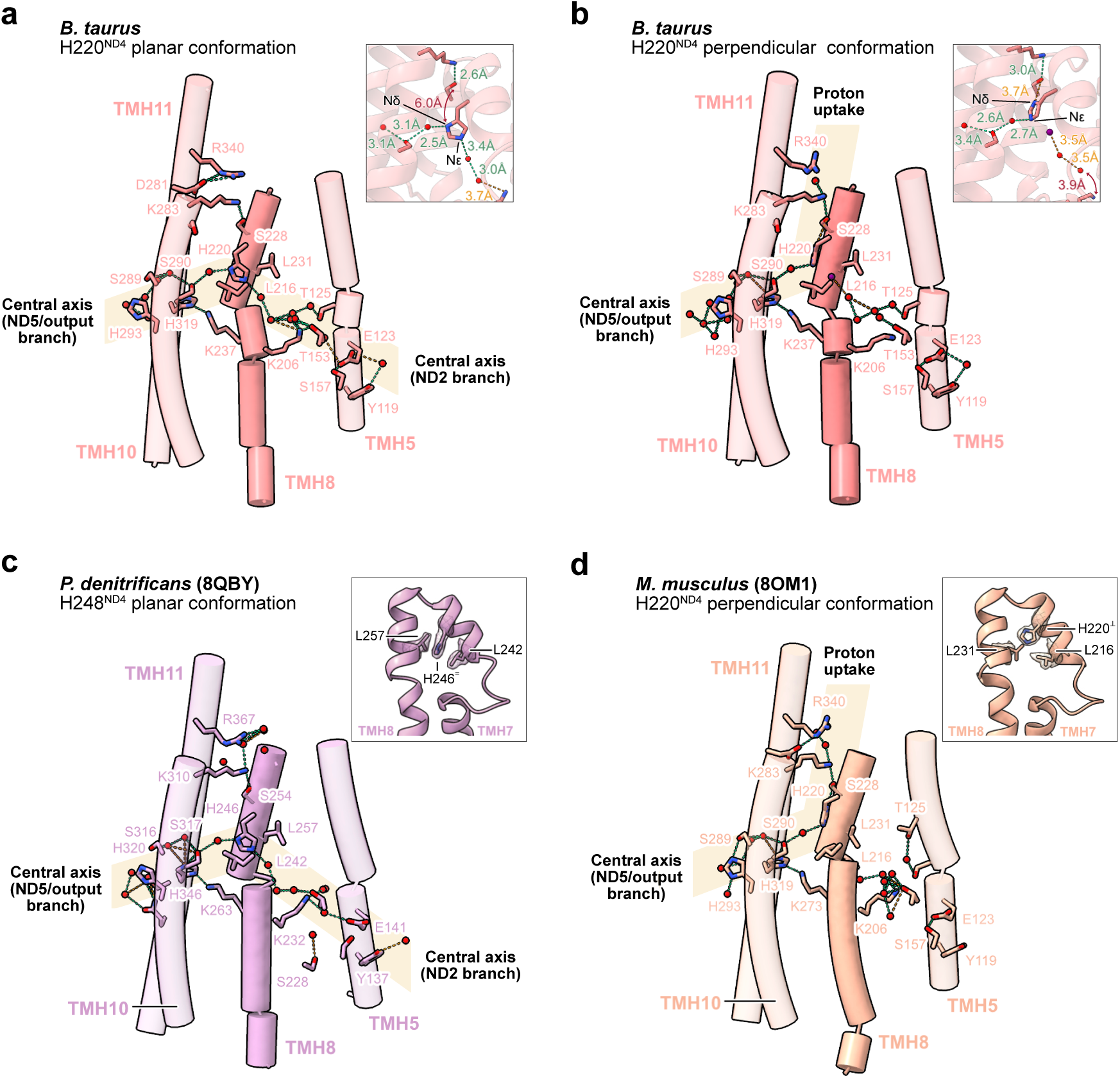
Rotamer changes in H220^ND4^ define proton transfer routes in ND4. (**a**) The planar position of H220 connects the ND2 and ND5/output branches of the central axis. (**b**) The perpendicular position of H220 connects the proton uptake channel to the ND5/output branch of the central axis. Inset shows hydrogen bonding interactions made by H220. Though both conformations can be modelled at each pH, only the highest confidence models with the greatest hydrogen bond connectivity are shown (the planar conformation from the pH 9.2 dataset and the perpendicular conformation from the pH 6.0 dataset; see Extended Data Fig. 6c for the planar conformation at pH 6.0 and perpendicular conformation at pH 9.2). (**c**) The ND4 subunit of complex I from *P. denitrificans* (PDB:8QBY) adopts a singular planar conformation. Inset shows the cryo-EM density for H246 positioned between L257 and L242. (**d**) The ND4 subunit of complex I from *Mus musculus* (PDB: 8OM1), adopts a singular perpendicular position. Inset shows the cryo-EM density for H220 rotated away from L231 and L216. Hydrogen bonding interactions are shown as green (<3.4 Å) and orange (3.4 – 3.7 Å) dashes and water molecules as red spheres, with the ambiguous water in the pH 6.0 structure (see Fig. 1d) in purple. Regions of continuous connectivity are highlighted in pale orange

In the planar conformation, H220 forms a junction between the two branches of the central axis. Its imidazole ring is sandwiched between two well-resolved conserved leucines (TMH7-L216 and TMH8-L231) that stabilize it through methyl CH-π interactions (Fig. 3a). H220-Nδ forms a water-mediated connection to S290, connecting H220 to the ND5/output branch of the central axis. Concurrently, a chain of three waters extends from H248-Nε to T153, which further connects to the K206-E123 ion-pair on the ND2 branch. These interactions are closely conserved in complex I from *P. denitrificans*^13^ (Fig. 3c) and bovine CI-PLs^41^ (Extended Data Fig. 6a), which both resolve H220 in a singular planar conformation, lending confidence to the hydrogen-bonding connections in Fig. 3a and identifying the planar rotamer as a bridge for lateral proton transfer across the central axis.

In the perpendicular conformation, H220 forms a junction between the proton-uptake pathway and the ND5/output branch of the central axis. Its imidazole ring moves away from the tandem-leucine motif and towards the putative uptake pathway (Fig. 3b), where H220-Nδ forms a hydrogen bond to S228, establishing a continuous connection to K283 at the top of the pathway. H220-Nε faces the ND5/output branch of the central axis, where a single water (present in both conformations) shifts its hydrogen-bonding partner from Nδ to Nε, maintaining its connectivity to S290. This configuration is homogeneously represented in structures from *M. musculus*^8^ (Fig. 3d) and *E. coli*^12^ (Extended Data Fig. 6b), though the structures diverge at L231, which is poorly resolved in mouse and substituted as *Ec*-I262 in *E. coli*. Further variation in *E. coli* sees S228 in the proposed proton-uptake channel replaced by an aspartate (*Ec*-D258).

Our structures suggest that the L216-L231 pair ‘corrals’ H220, helping to direct it during catalysis. The functional significance is underlined by simulations that indicated L216 gates hydration of the proposed proton-uptake channel^31,49^, and by the L216A mutation in *P. denitrificans* (*Pd*-L242A) that decreases the rate of catalysis by ∼90%^27^. The *Pd*-L242A mutation also drastically alters the pH-dependence of catalysis, shifting the pH optimum from ∼7.0 to ∼8.0, a unique effect among available ND4 central axis mutations (Extended Data Fig. 7). Together, these findings identify H220 as a proton-transfer junction that, analogous to H248^ND5^, switches the connectivity between the central axis and matrix-uptake pathways to coordinate vectorial proton transport.

### Coupling lysine protonation to histidine gating

Our structures define a clear protonation-dependent switch at the H248^ND5^ junction, but the physical trigger for the rearrangement is unclear. Given the canonical histidine pKa of ∼6.0 [neutral to HisH^+^] falls in our experimental range, we first reasoned that H248^ND5^ itself might change protonation state to induce the switch. However, constant-pH molecular dynamics (CpHMD) simulations, which account for the global membrane environment as well as local hydration and sidechain dynamics, indicated that the buried H248^ND5^ remains neutral at both pHs (Fig. 4a) (protonated on either Nδ or Nε, 62 ± 18% on Nε at pH 6, 72 ± 20% at pH 9.2). Instead, they identified protonation changes at the top of the proposed proton-uptake pathway, where K299 and K300 appear fully protonated at pH 6.0 (combined charge +2) but partially deprotonated (+1.3) at pH 9.2. Additionally, E145, in the ion-pair where the central axis enters from ND4, switches from a mixed state at pH 6.0 (–0.45) to deprotonated at pH 9.2 (–1). These results suggest the H248^ND5^ conformation is not set by the protonation state of the histidine itself, but by the states of titratable residues on pathways connected to it.

**Figure 4.**
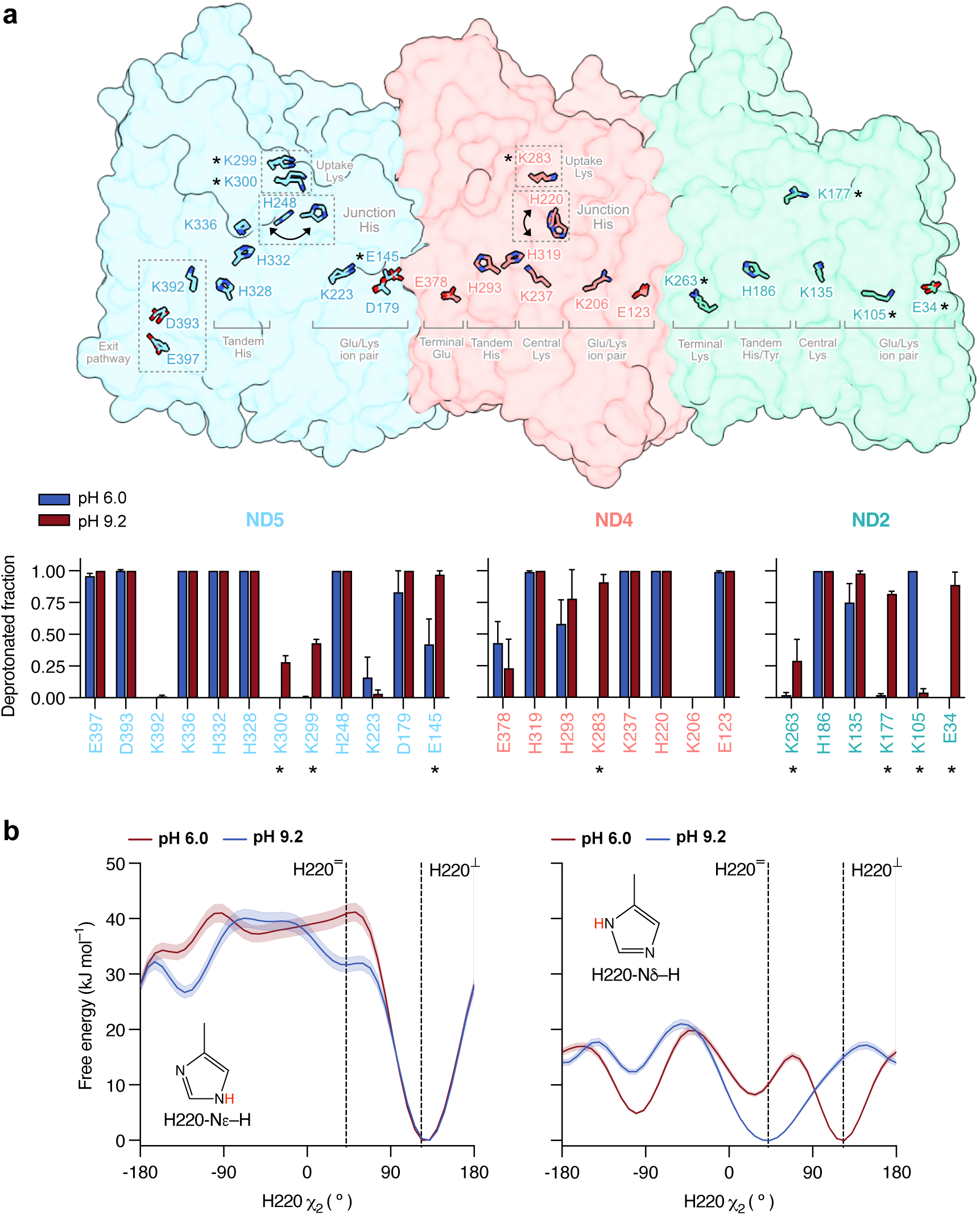
Protonation-state changes in the antiporter-like subunits of complex I. **(a)** The surface of the membrane subunits ND2, ND4 and ND5 are shown and the positions of titratable residues at pH 6.0 and 9.2 are indicated. The movement of the junction histidine residues is highlighted by arrows. The protonation states obtained from CpHMD simulations are shown in the bar charts and residues that show a change of >25% in their protonation state between the pH values are indicated with an asterisk. Densities for asterisked residues are shown in Extended Data Fig. 8. (**b**) Free energy profiles calculated for torsion of the H220^ND4^ χ_2_ dihedral in both tautomeric states at pH 6.0 and 9.2. Neutral forms of H220^ND4^ protonated at Nε (HSE, left) and Nδ (HSD, right) are shown. χ_2_ angles that represent the planar (H220^=^, 42°) and perpendicular (H220^⊥^, 123°) states are indicated. Error bars represent the free energy uncertainty. For reference, RT ∼2.5 kJ mol^−1^. See Supplementary Fig. 7 for data on the convergence of the simulation.

Similar behaviour is observed in ND4, where CpHMD indicates that H220 also remains neutral at both pHs (Fig. 4a). K283, at the top of the proposed proton-uptake pathway deprotonates cleanly (+1 to 0) between pH 6.0 and 9.2, but no change is predicted at the ion pair (E123-K206). However, in ND4 the perpendicular and planar states are themselves less cleanly interconverted by pH, so we performed metadynamics simulations with enhanced sampling of the χ1 and χ2 dihedrals of the H220 sidechain to investigate the energy landscape. Using CpHMD-determined protonation states for the surrounding residues, we found the conformation of H220 is highly sensitive to its tautomeric state (Fig. 4b). When Nε is protonated [HSE], the perpendicular conformation is strongly favoured regardless of pH. Conversely, both forms are populated when Nδ is protonated [HSD], with the perpendicular conformation favoured at pH 6.0 and the planar conformation at pH 9.2. Our data suggest the uptake-pathway lysine influences the tautomeric and conformational landscape of the junction histidine, which functions as a switch that controls the proton-transfer connectivity.

The CpHMD analysis also highlights protonation changes in ND2 (Fig. 4b), which lacks a junction His and shows no material structural changes (Extended Data Fig. 9). K177^ND2^, in the same position as K300^ND5^ and K283^ND4^ at the top of the proposed proton-uptake channel, switches protonation state, while protonation inversion of the E34-K105 ion-pair (–/+ at pH 6.0, 0/0 at pH 9.2) may reflect differential modelling of nearby water molecules.

Although our results do not establish a cause-effect relationship, they suggest the matrix-facing lysines in ND4 and ND5 may act as local control points, triggering changes at the junction histidines to configure the connectivity of proton-transfer pathways.

## Discussion

The chemical potentials of the protons on each side of the energy-transducing membrane are important contributors to the thermodynamic driving force for complex I catalysis, which dictates its direction and efficiency. Although altering the bulk pH cannot create a potential difference across the membrane domain of the isolated enzyme, changing it from 6.0 to 9.2 changes the proton chemical potential around the enzyme by ∼180 mV, equivalent to the proton-motive force *in vivo*. Here we aimed to use the bulk pH to shift the protonation states of catalytically crucial residues, and thereby to induce new conformational states to inspire insights into mechanism.

Given the significant energetic shift we impose, it is surprising that only two structural changes are observed in the substantial proton-translocating membrane domain, even at 2.2–2.3 Å resolution. Until now, conformational changes within a single species of complex I, including those determined under conditions designed to stimulate turnover^11,12,15,50^, have been limited to the substrate-binding regions, or linked to the regulatory interconversion of closed and open states that involves the Q-site and E-channel. In contrast, the changes we identify here are in the antiporter-like subunits, where altered proton energetics likely act most directly. The number of structural changes may be limited because most catalytically active residues in the membrane domain are buried, only connected to the bulk pH by confined hydrogen-bonding networks that accommodate protonation changes through subtle rearrangements in connectivity and hydrogen-bond directionality and resist large changes in proton occupancy.

### H248 as a master-switch in subunit ND5

Our high-resolution structures of mammalian complex I reveal a conformational switch at H248^ND5^ that controls proton-transfer connectivity at a trigonal junction between the ND4-upstream and output-downstream branches of the central axis and the proposed proton-uptake pathway. The substantial helical rearrangement is unmatched in ND4 or ND2, suggesting it as a ‘master-switch’ in the proton-pumping mechanism. Although our structures provide the first direct experimental evidence for such a switch in complex I, both conformations have been observed separately in structures from different species^12^ and molecular dynamics simulations registered mobility in this region^28^. Studies of the homologous MrpA in the evolutionary-linked MRP-type antiporters identified alternative conformations for H248 (but without resolving two discrete states)^29^. Subsequently, by combining structural, mutational and computational analyses on MrpA, a histidine-switching proton-transfer mechanism was proposed, and three lysines implicated in controlling the switch during catalysis^30^. Protonated K299 (uptake) was suggested to stabilise the contracted state and protonated K336 (output) the expanded state, whereas our data on complex I suggest that protonation of the K299–K300 pair (uptake) stabilises the expanded state and K336 (output) remains deprotonated. However, our structures are of resting states, not intermediates, and may be kinetically trapped: the high-pH structure may be stuck in the contracted state because deprotonated K299 cannot deliver a proton, and the low-pH structure in the expanded state because contraction is redox-coupled or requires catalytic steps not accessible under our conditions. In any case, our structures define distinct, interconvertible states that switch the proton-transfer connectivity at a key pathway junction in the membrane domain of complex I.

Fig. 5a illustrates a model for gated proton transfer through ND5, similar to the MrpA-based mechanism proposed previously^30^. H248^ND5^ switches between the contracted and expanded conformations while cycling between the singly (HSE) and doubly (HSP) protonated states. In the contracted HSE state, Nδ is positioned to receive a proton from K299^ND5^ via the proton-uptake pathway. Protonation generates the HSP state, which transitions to the expanded conformation, connecting Nδ to the proton-output pathway to donate its proton to K336^ND5^. The neutral HSE then switches back to the contracted conformation, completing the cycle. However, this cycle accounts only for local translocation, it does not explain how proton pumping is redox coupled or reverse proton leak prevented. One possibility is that signals from redox catalysis are transmitted via the central axis to H248^ND5^-Nε, which is connected to the ND4 branch in the contracted state. Notably, neither conformation links H248^ND5^-Nε to the output pathway so an additional (unknown) state is needed if protons are to be transferred from ND4 into the IMS by H248^ND5^ and the ND5 output pathway.

**Fig. 5.**
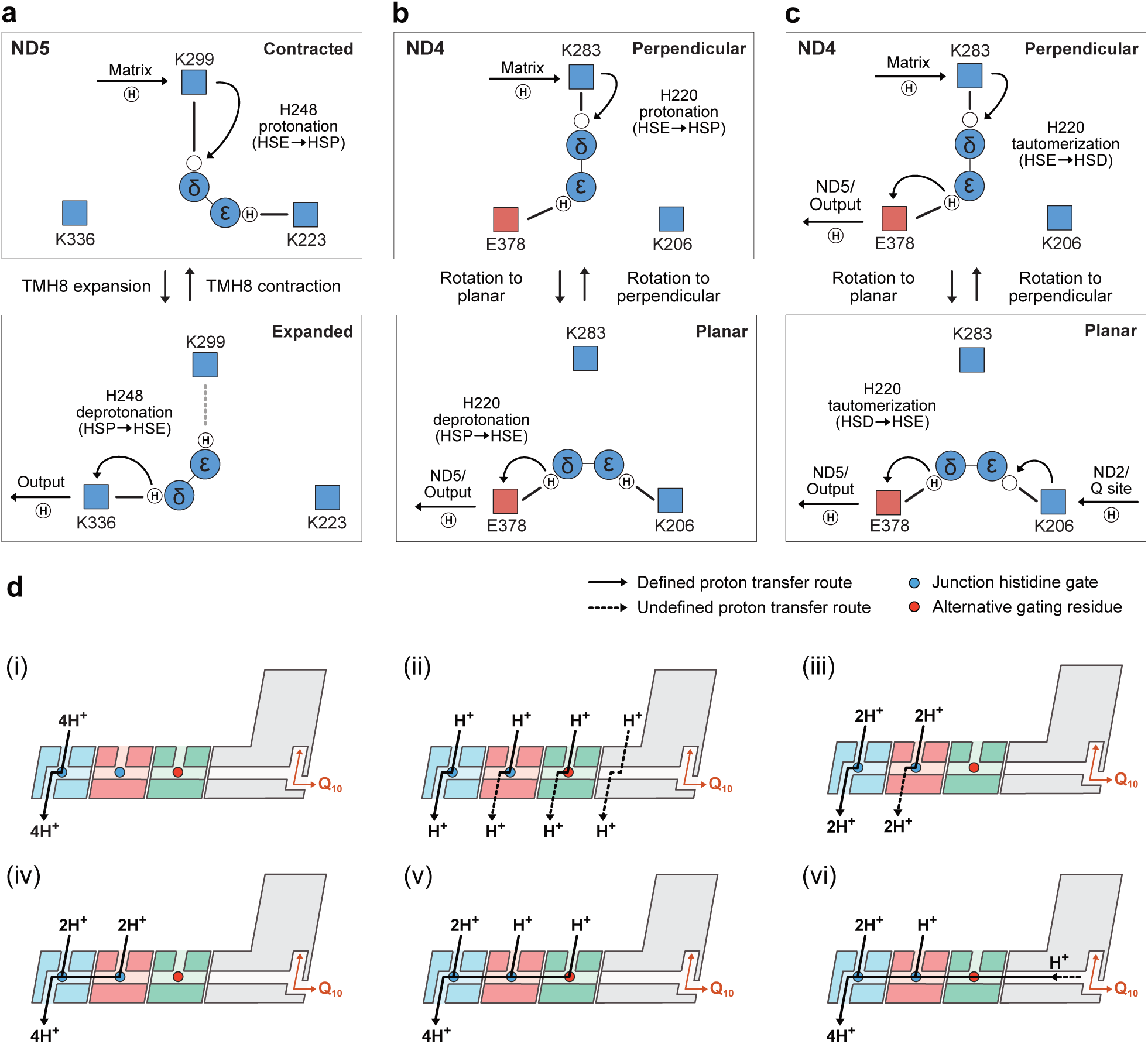
Gated proton transfer in the membrane domain of complex. **I.** (**a**) Conceptual scheme for conformationally gated proton transfer from the uptake to the output pathway by the master switch in ND5. The three experimentally determined pathways from H248 are represented by: K299, proton uptake; K223, ND4 branch of central axis; K336, proton output. Well defined pathway connections are indicated by solid black lines, with a dotted grey line indicating an ambiguous connection. A proton is taken up from the matrix via K299 in the contracted state and transferred to H248-Nδ. The resulting HisH^+^ (HSP) transitions to the expanded state and H248-Nδ donates the proton to the output pathway via K336. H248 returns to the contracted state. (**b**) Conceptual scheme for rotamer gated proton transfer from the uptake to the ND5/output pathway in ND4. The three experimentally determined pathways from H220 are represented by: K283, proton uptake; K206, ND2 branch of central axis; E378, proton output. A proton is taken up from the matrix via K283 in the perpendicular state and transferred to H248-Nδ. The resulting HisH^+^ (HSP) rotates into the planar state and H248-Nδ donates the proton to the ND5/output pathway via E378. H220 rotates back to the perpendicular state. (**c**) Conceptual scheme for tautomeric gating of a double-proton transfer reaction in ND4. In the perpendicular state, a proton is taken up from the matrix and transferred to H220-Nδ and the proton on H220-Nε is donated to the ND5/output pathway. H220 rotates into the planar state, where a proton is taken from the ND2 branch of the central axis H220-Nε via K206 and the proton on H220-Nδ is donated to the ND5/output pathway. H220 rotates back to the perpendicular state. **(d)** Prospective models of proton translocation in complex I. The ND5, ND4 and ND2 subunits are shown in blue, pink and green respectively. Histidine switches are indicated by a blue circle, with alternative residues (Thr, Tyr), shown in red. Solid lines represent structurally defined proton transfer routes, while dashed lines represent suggested routes of proton transfer that have not been identified in structural data.

### Tautomeric gating and the ND4 clockwork mechanism

In ND4, H220 similarly forms a trigonal junction between the ND2-upstream and the ND5/output-downstream branches of the central axis (i.e. protons may be output from ND4 or be transferred to ND5) and the proposed proton-uptake pathway. A simple proton-transfer cycle analogous to that described for ND5 can be envisaged in which H220-Nδ transfers protons (from uptake to ND5/output) by rotating between the perpendicular and planar conformations (Fig. 5b). However, while the distances between the connections to H248^ND5^ in the contracted and expanded conformations require the doubly protonated HSP to ferry protons across the gaps, the distances are shorter in ND4 and both Nδ and Nε of H220^ND4^ remain connected in both states. A tautomeric gating mechanism, which does not require HSP, can thus be constructed (Fig. 5c). In the perpendicular HSE state (favoured in our simulations, Fig. 4b), protonation of H220-Nδ from the uptake pathway is coupled to deprotonation of H220-Nε to the ND5/output branch of the central axis. The resulting HSD tautomer is unstable in the perpendicular conformation and rotates to the planar conformation, bridging the two branches of the central axis. H220-Nε collects a proton from the ND2 branch while H220-Nδ delivers a proton to the ND5/output branch, restoring the HSE tautomer, which returns to the perpendicular conformation completing the cycle. This ‘clockwork’ mechanism shuttles the protons from two incoming pathways alternately into a single onward pathway. Although a powerful concept for ND4, extending this mechanism to ND5 would require additional complexity to either exchange the histidine tautomer (for example, transiently passing a proton from Nε to the uptake pathway then accepting it back onto Nδ), or to connect Nε to the output pathway in a so-far unobserved intermediate.

### Towards a unified proton-wiring diagram

Finally, we consider how the gating mechanisms in Fig. 5a-c could combine to define the proton-wiring diagram for complex I catalysis (Fig. 5d). If the H248^ND5^ master switch operates only through the structurally defined cycle (Fig. 5a), then ND5 functions independently and only transports protons from the matrix to the IMS. The key question then becomes how many protons follow this route during each NADH oxidation cycle. All four protons could be transferred by ND5 (Fig. 5d-i), consistent with it containing the only structurally defined proton-output channel. However, it is unclear how four identical proton-pumping cycles could be driven sequentially by a single redox cycle and the H220 gate in ND4 suggests a more integrated circuit. If H220^ND4^ functions similarly to H248^ND5^ (Fig. 5b), then ND4 may also function as a self-contained transporter and a series of transporter modules become possible. The four-module model, in which ND5, ND4, ND2 and a fourth putative module in the proximal membrane domain each translocate a proton (Fig. 5d-ii), requires each subunit to possess defined gating mechanisms and proton uptake and output pathways. However, gating mechanisms in ND2 and the fourth-module are currently unknown, along with a fourth-module uptake channel, and ND4, ND2, and fourth-module output channels. Alternatively, ND5 and ND4 might each contribute two protons per cycle (Fig. 5d-iii), restricting the challenge to lack of evidence for an output pathway from ND4. If one does exist then it likely begins at TMH12-E378^ND4^, adjacent to the ND5 ion pair, tentatively suggesting that the master switch may coordinate its activity across the ND4/5 interface, via H248^ND5^-Nε and the central axis.

An alternative class of mechanisms invokes a ‘clockwork’ role for the H248^ND5^ master switch in which both Nδ and Nε participate, allowing protons to be routed from upstream subunits and expelled via the structurally defined output pathway. In this ND5 clockwork mechanism, H248^ND5^ alternately shuttles an uptake-pathway proton on Nδ and an ND4-upstream central-axis proton on Nε to the same K336-mediated ND5-output pathway. The challenge to this mechanism is lack of structural evidence for a state that connects Nε to the output pathway. In the simplest case (Fig. 5d-iv), ND4 and ND5 each take up two protons, the central axis transfers the protons from ND4 to ND5, and the clockwork master switch controls output of all four through a universal gating mechanism. This simplest case, however, does not use the ND4 clockwork mechanism, which is supported structurally. In a related model, ND2 and ND4 each take up one proton; they are combined by the ND4 clockwork mechanism, then combined with two further protons taken up by ND5, to all exit from ND5 (Fig 5d-v). In a further variation, a single proton taken up by ND4 combines instead with a ‘driving’ proton transferred across ND2 from redox catalysis (Fig. 5d-iv), as a speculative suggestion that links the redox and proton-pumping cycles.

In summary, gating mechanisms in the membrane domain of complex I impose directionality on proton-transfer mechanisms and eliminate uncoupling short circuits by gating connections to proton uptake and output pathways. The master switch at H248^ND5^ and the clockwork mechanism at H220^ND4^ prevent ungated (unregulated) connections and likely represent key control points in catalysis. How they are controlled and driven by the redox reaction, and whether they act as independent control points or within a single integrated multi-subunit module define key questions at the heart of the complex I mechanism that will form the focus of future studies.

## Online Methods

### Preparation of native membrane systems

*Bos taurus* (bovine) heart mitochondria and mitochondrial membranes were prepared using established protocols^42^. Sub-bacterial particles (SBPs) were prepared from variants of the *Pd*-Nqo5^His6^ strain^51^ by growing *P. denitrificans* aerobically to the mid-exponential phase in LB media. SBPs were then prepared by osmotic lysis as described previously^27,51^.

### Preparation of *B. taurus* complex I and complex I-containing proteoliposomes

Mitochondrial membranes were solubilized with DDM and complex I purified using anion exchange and size exclusion chromatography as described previously^42,43^. For cryo-EM samples, complex I-containing anion-exchange fractions were pooled and concentrated then injected onto a Superose™ 6 Increase 5/150 GL column (Cytiva) pre-equilibrated in Tris-MES buffer (15 mM Tris, 15 mM MES, 150 mM NaCl, 0.05% DDM) corrected to pH 6.0 or 9.2 at 4 °C. Protein concentrations were determined using the bicinchoninic acid (BCA) assay kit (Pierce).^52^ Complex I for proteoliposome (PL) studies was purified in LMNG^41^ before a sodium cholate-mediated reconstitution into preformed Q_10_-supplemented liposomes (8:1:1 (by weight), 1,2-dioleoyl-sn-glycero-3-phosphocholine (DOPC), 1,2-dioleoyl-sn-glycero-3-phosphoethanolamine (DOPE), and 18:1 cardiolipin (TOCL)) as described previously^41^. Following ultracentrifugation, CI-PLs were resuspended in a minimum volume of 10 mM MOPS (pH 7.5), 50 mM KCl and the concentration of complex I determined using the NADH:APAD^+^ oxidoreduction assay (see below). CI-PLs were diluted to 100 µg mL^−1^ in CHB buffer (20 mM citric acid, 30 mM HEPES, 40 mM boric acid) at the desired pH for a minimum of 1 hour on ice to equilibrate the pH before activity measurements.

### Complex I activity measurements

All complex I activities were measured spectrophotometrically at 32 °C using a SpectraMax® ABS Plus 96-well microplate reader (Molecular Devices). Rates of NADH:decylubiquinone (DQ) oxidoreduction were determined in 20 mM Tris-HCl (pH 7.5 @ 32°C) with 0.5 µg mL^−1^ complex I, 0.15% soybean asolectin (Avanti polar lipids), 0.15% CHAPS and 200 µM DQ. Catalysis was initiated by addition of 200 µM NADH and oxidation of NADH measured at 340–380 nm (ε_340–380_ = 4.81 mM^−1^ cm^−1^).

The concentration and orientation of complex I in CI-PLs were determined from the rate of NADH:3-acetylpyridine adenine dinucleotide (APAD^+^) oxidoreduction at the complex I flavin site. NADH:APAD^+^ rates were determined at 32 °C in 10 mM MOPS (pH 7.5), 50 mM KCl, 500 µM APAD^+^ and 1 µM piericidin A. Catalysis was initiated with 100 µM NADH and reduction of APAD^+^ followed at 450–400 nm (ε_450–400_= 3.16 mM^−1^ cm^−1^). The complex I concentration was determined by comparison to the rate of a LMNG-solubilized complex I standard. The proportion of inward and outward-facing complex I (CI-out) was determined by adding 15 µg mL^−1^ alamethicin to permeabilize the membranes to NADH and APAD^+^.

For both CI-PLs and native membranes, NADH:O_2_ oxidoreduction rates were measured by monitoring NADH oxidation, with the alternative quinol oxidase (AOX) from *Trypanosoma brucei brucei* added to regenerate oxidized ubiquinone^52,53^. In *B. taurus* mitochondrial membranes, canonical respiratory chain activity was inhibited with 400 µM NaCN and AOX was added at 1 mg per mg (total membrane protein) to focus the measurements on complex I activity only. Catalysis was initiated by adding 200 µM NADH. For SBPs from *P. denitrificans*, AOX was added at 0.5 mg per mg (total membrane protein), with 1 µM antimycin and 400 µM NaCN to inhibit complexes III and IV. To avoid non-specific NADH oxidation by NDH2, the complex I specific deamino-NADH (dNADH) was used for SBPs. For maximum catalysis, the proton motive force was dissipated using 1 µg mL^−1^ gramicidin D (GramD). Rates of NADH oxidation in proteoliposomes were measured with 0.5 µg/mL CI-out, 5 µg/mL AOX and 200 µM NADH (± 0.5 µg/mL GramD for uncoupling). In all cases, pH dependent activities were measured in CHB buffer across a pH 4 – 10 range (set at 32 °C). For pH stability measurements, membranes, SBPs and CI-PLs were incubated for 1 hour at 4 °C in CHB buffer across a pH 4 – 10 range (set at 4 °C) prior to determination of the activity at pH 7.5.

### Biochemical determination of resting states

The relative rate of NADH oxidation following treatment with *N*-ethylmaleimide (NEM) was used to assess open/D and closed/A state proportions. CI-PLs were diluted to 100 µg mL^−1^ in CHB buffer at the desired pH for a minimum of 1 hour on ice to equilibrate the pH. Samples were then labelled with 1 mM NEM for 30 min before determination of the NADH:O_2_ activity. Conversion of CI-PLs to an all-open/D state^41^ was achieved by the standard deactivation procedure of 20-min incubation at 37 °C. To avoid effects of differential stability at different pH values, thermal deactivation was performed at pH 7.5 prior to incubation at the experimental pH. At pH 7.5, complex I activity was fully recoverable following deactivation (but this was not the case at pH 6.0, Extended Data Fig. 2d).

### Cryo-EM grid preparation and data acquisition

UltrAuFoil® gold grids (R 0.6/1, Quantifoil® Micro Tools GmbH) were glow discharged at 30 mA for 90 s then derivatized with a PEG-thiol compound, SPT0011P6 (SensoPath Technologies, Inc.), as described previously^40,54^. 3 µL of 6 mg mL^−1^ complex I was applied to each grid at 4 °C and 100% relative humidity, blotted for 10 s at blot force -10, then plunge frozen using an FEI Mark IV Vitrobot (Thermo Fisher). Grids were screened for ice quality and particle distribution using a Talos™ Arctica™ microscope (Thermo Fisher) at the Department of Biochemistry, University of Cambridge.

The pH 6.0 dataset was collected at the UK National Electron Bio-Imaging Centre (eBIC) and the pH 9.2 dataset was collected at the Department of Biochemistry, University of Cambridge. Each dataset was collected on a 300 keV Titan Krios™ microscope (Thermo Fisher) with a K3® detector and Quantum® energy filter (Gatan) with a 20 eV slit width operating in counting mode, with 100 µm objective and 70 µm C2 apertures and using EPU software (Thermo Fisher). Datasets were collected with Aberration-free Image Shift (AFIS) and Fringe-free Imaging (FFI) with a 5 s delay after stage shift and 1 s delay after image shift and defocus ranges of -2.3 µm to -0.9 µm (0.1 µm increments). Super-resolution movies were collected at 0.532 (pH 6.0) or 0.533 (pH 9.2) Å/px (81000× nominal magnification) then 2× Fourier binned to 1.064 (pH 6.0) or 1.066 (pH 9.2) Å/px. Autofocus programs were run every 10 µm. The pH 6.0 dataset was acquired with one 2.80 (pH 6.0 or 2.67 (pH 9.2) s exposure per hole with a beam diameter of 1.30 (pH 6.0) or 1.35 (pH 9.2) µm (40 frames), and 14.6 (pH 6.0) or 15.2 (pH 9.2) electrons Å^-2^ s^-1^ (total dose 40.9 electrons Å^-2^).

### Cryo-EM image processing

Both datasets were processed using matching pipelines of both RELION v4.0 beta^55^ and CryoSPARC v3.3.2^56^ (Supplementary Fig. 1). 21,685 (pH 6.0) or 12,549 (pH 9.2) curated micrographs were Fourier cropped by 2× then corrected for beam-induced motion using RELION’s implementation of MOTIONCOR2 with 10×10 patches and an amplitude contrast proportion of 0.1, and contrast transfer function (CTF) estimated using CTFFIND-4.1 with a ResMax of 4 Å. Acceptable micrographs with -0.05 ≤ rlnCtfFigureOfMerit ≤ 3 and rlnMaxResolution ≤ 6 Å were retained, and micrographs with poor ice thickness or quality were manually removed, leaving 14,680 (pH 6.0) or 8,869 (pH 9.2) micrographs. RELION’s Autopicker was used with a previous 3D reference map of bovine complex I (7QSN^22^), yielding 1,446,706 (pH 6.0) or 1,056,616 (pH 9.2) initial particles, and particles were then extracted at 2.394 (pH 6.0) or 2.3985 (pH 9.2) Å/px, respectively. For each dataset, an initial round of 3D classification (six classes) was used to remove junk particles, with three complex I classes retained. A second round (five classes) was performed and three complex I classes retained, each having characteristics of the global states. Particles were combined and then re-extracted back to 1.064 (pH 6.0) and 1.066 (pH 9.2) Å/px.

The particle stacks were subjected to alignment-free 3D classification in CryoSPARC (four classes), providing volumes for use as references for subsequent heterogeneous refinement, yielding three complex I classes for each dataset. A mixed open/slack class from the pH 6.0 dataset was separated by a further round of heterogeneous refinement. Homogeneous refinement with per-particle defocus refinement and global CTF refinement was then performed with the particle stacks imported into RELION. To further improve per-particle motion, RELION’s Bayesian polishing^57^ was trained using a CTF-refined open/D class from the same dataset and each class was polished and re-extracted at 0.798 (pH 6.0) and 0.800 (pH 9.2) Å/px, then refined. The polished particle stacks were imported back into CryoSPARC for a final round of homogeneous refinement with per-particle defocus refinement and global CTF refinement. The final particle numbers were 799,398 (247,586 closed/A, 301,984 open/D, 249,828 slack) for the pH 6.0 dataset and 718,227 (249,186 closed/A, 262,659 open/D, 206,422 slack) for the pH 9.2 dataset (Supplementary Figs. 2 and 3). To further validate the proportions closed/A particles, the focus-revert-classify approach was employed as described previously^58^ (Supplementary Fig. 4). Briefly, the polished and refined particle stacks from RELION for each class were joined, and a randomized 10% subset taken and refined. A soft mask was created for membrane and peripheral domains from each dataset in RELION and each subset focus refined using a peripheral domain mask and classified with a membrane domain mask, with class ratios then compared. Following import back to RELION using the csparc2star.py script in the pyem package [10.5281/zenodo.3576630]^59^, particle stacks underwent iterative rounds of CTF refinement for all global and local parameters (initially starting with correction for anisotropic magnification) followed by 3D refinement.

All masks were created using the molmap command in ChimeraX-1.8^60^ from previous bovine models (7QSL and 7QSN^22^), lowpass filtered to 17 Å, with a soft edge of 10 px. The closed/A and slack consensus maps were sharpened in RELION with B-factor estimation at a minimum resolution of 10 Å. Local half map-based sharpening was applied to the closed/A and slack maps using *phenix.auto_sharpen* in Phenix-1.20.1^61^, with maps clipped to include only the region of interest using *phenix.map_box* and coordinates from a previous model (7QSN^22^). To generate the composite open/D maps, soft regional masks were dilated and padded in CryoSPARC (Supplementary Fig. 5). The masks were subsequently inverted and used in a particle subtraction and local refinements then undertaken (Supplementary Fig. 6). The B-factor-sharpened focused maps were combined in Phenix-1.20.1 with *phenix.combine_focused_maps* using a previous open/D state model (7QSN^22^) for rigid body fitting into each B-factor-sharpened focused map and original consensus map. The composite map was used to aid model building of the open/D state and for the first round of *phenix.refine* with all subsequent refinements using the global consensus maps. Local resolutions were estimated using CryoSPARC with masks from global or local refinements and visualized on ChimeraX.

### Model building, refinement, validation and analysis

Hydrogen atoms were added to bovine complex I models from a previous study (7QSL for the closed/A states and 7QSN for the open/D states^22^) using *phenix.readyset*, then rigid body-fitted into the pH 6.0 and 9.2 maps using ChimeraX, and all-atom and chain-refined using Coot 0.9.8.92^62^. Residue density misfits were also checked using the model-map atom resolvability *Q-score*^63^ implementation in ChimeraX^60^. Existing waters were manually inspected and adjusted and additional waters placed in clear density peaks using the Find Waters tool in Coot. All newly added waters were manually inspected. Multiple rounds of *phenix.refine* were undertaken with all refinements of the open/D models into global consensus maps, with hydrogens removed from each model before the final round. Model statistics were generated using the Phenix implementation of MolProbity v4.4 during the final round of *phenix.refine* with model-to-map FSC curve data using *phenix.mtriage* (Supplementary Tables 1 and 2 and Supplementary Fig. 3). Hydrogen bonds were identified with the hbonds command in ChimeraX^60^ with the default settings (max 3.4 Å) and at a higher tolerance for distance (3.7 Å).

### Molecular dynamics simulations

Two simulation models based on the pH 6.0 and 9.2 closed/A structures determined here were built, retaining only subunits ND5, ND4, ND2, and ND4L for efficiency. Each subcomplex was embedded in a previously equilibrated bilayer composed of 90 phosphatidylcholines, 80 phosphatidylethanolamines, 26 cardiolipins (di-anionic), and four oxidized Q_10_ to mimic the inner mitochondrial membrane. Phospholipids were built with linoleoyl (L, 18:2) acyl chains. Supplementary Table 4 lists the composition and total time of all the simulations performed.

All simulations were conducted at constant temperature (310 K, with the Bussi thermostat) and pressure (1 atm, with the Parrinello-Rahman barostat, except the initial equilibration which used the Berendsen barostat), and a time step of 2 fs. Long-range electrostatics were treated by the Particle Mesh Ewald method. Interactions of the protein, lipids, and ions were described using the all-atom CHARMM36m force field ^64^. Water was represented by the standard TIP3P model. Calibrated parameters were used for Q_10_^65,66^. Simulations were run with GROMACS version 2024^67^, except when noted. For the initial set-up, and membrane and solvent equilibration, internal water molecules modelled in the cryo-EM structures were retained. Protonation states of all residues were adjusted to neutral pH, except for E144^ND4^ and E34^ND4L^, which were always protonated due to being exposed to the membrane from the simulated subcomplex. Harmonic restraints of 1000 kJ mol^−1^ nm^−2^ were applied to all protein heavy atoms. Membrane thickness and area per lipid, and internal protein hydration were verified to be stable during the last 80 ns of simulation.

To assign the relevant side-chain protonation states, constant-pH molecular dynamics (CpHMD) simulations were carried out with GROMACS version 2021-beta1 (https://www.gitlab.com/gromacs-constantph)^68^. Supplementary Table 5 lists the 42 residues listed titrated. Some λ-bias barriers were changed from the default 7.5 kJ mol^−1^ to minimize sampling of unrepresentative partial protonation states. CpHMD-specific modifications of the CHARMM bonded parameters for titratable residues were used. Buffer particles were added to account for charge fluctuations and maintain a neutral system^69^. CpHMD simulations were run in triplicate for each model at its respective pH (Supplementary Table 4). The first (longer) replicate at each pH started after solvent equilibration, with protein restraints reduced to zero in two steps until 20 ns. Subsequent replicates started from the 200 and 300 ns configurations of the first replicate, with randomized initial protonation states of titrated residues. Protonation states (λ values) were recorded every 2 ps and protonation fractions computed as time-averaged values, after removing the initial 30 ns for additional equilibration, with protonated or deprotonated states cut off at λ < 0.2 and > 0.8, respectively. Reported fractions (Fig. 4) are the mean ± standard error from the three replicates.

To probe the relative stability of H220^ND4^ conformers, well-tempered metadynamics^70^ with fixed protonation states were performed for each model, with H220 set to neutral Nε-protonated (HSE) or Nδ-protonated (HSD) (Supplementary Table 4). These simulations started from the configuration at 100 ns of the first CpHMD replicate. Protonation states for each titrated residue were set to the most highly populated states from the CpHMD analyses (Supplementary Table 5), except for K283^ND4^ which was deprotonated in the pH 9 model. Interactions were described using the original CHARMM36m force field (without the CpHMD-specific modifications). Metadynamics were applied to the ND4 H220 χ1 (Cɑ-Cβ bond torsion) and χ2 (Cβ-Cγ) dihedrals, with Gaussians deposited every 500 time steps (1 ps), at an initial height of 0.6 kJ·mol^−1^, width of 0.02 units, and a bias factor of 15.0. Convergence within ±1.5 kJ mol^-1^ of free energy differences was reached after ∼150 ns (Supplementary Fig. 7). Metadynamics simulations were performed with the PLUMED plugin (version 2.9)^70^. Initial configurations for all MD simulations are available online (https://doi.org/10.5281/zenodo.18805727).

## Data and materials availability

The cryo-EM data generated in this study have been deposited in the electron microscopy databank (EMDB) and protein databank (PDB) The pH 6.0 datasets have the following accession codes: EMD-53490 and PDB-9R12 (pH 6.0 closed/A), EMD-53491 and PDB-9R13 (pH 6.0 open/D, consensus), EMD-53495 (pH 6.0 open/D, composite), EMD-53492 (pH 6.0 open/D, peripheral domain), EMD-53493 (pH 6.0 open/D, proximal membrane domain), EMD-53494 (pH 6.0 open/D, distal membrane domain), EMD-53496 (pH 6.0 slack). The pH 9.2 datasets have the following accession codes: EMD-53497 and PDB-9R14 (pH 9.2 closed/A), EMD-53498 and PDB-9R15 (pH 9.2 open/D, consensus), EMD-53502 (pH 9.2 open/D, composite), EMD-53499 (pH 9.2 open/D, peripheral domain), EMD-53500 (pH 9.2 open/D, proximal membrane domain), EMD-53501 (pH 9.2 open/D, distal membrane domain), EMD-53503 (pH 9.2 slack). Initial configurations for all MD simulations are available online (https://doi.org/10.5281/zenodo.18805727). All data needed to evaluate the conclusions in the paper are present in the paper and/or the Extended Data and Supplementary Information.

## Acknowledgments

We thank D. Chirgadze, S. Hardwick and L. Cooper (University of Cambridge Cryo-EM facility) for assistance with grid screening and the pH 9.2 data collection. Cryo-EM data for the pH 6.0 was recorded at the UK National Electron Bio-Imaging Centre at the Diamond Light Source, funded by the Wellcome Trust, MRC, and BBSRC. We thank A. J. Raine, E. E. Marcus, and A. J. Nelson (MRC MBU) for IT support and Hannah R. Bridges (MRC MBU) for assistance in collecting and processing cryo-EM data.

## Author contributions

JJW and JH conceived the project. WF prepared and characterised complex I for cryo-EM. WF performed all cryo-EM data processing and model building with support from DNG. WF and JJW performed structural analysis with input from DNG. JJW and MNC performed biochemical analyses at different pHs. GMA performed molecular simulations. JJW, RAW and JH interpreted the data and proposed mechanistic hypotheses. JH supervised the project with input from JJW. GMA and JH acquired funding. JJW and JH wrote the manuscript with input from all authors.

## Extended Data

**Extended Data Fig. 1.**
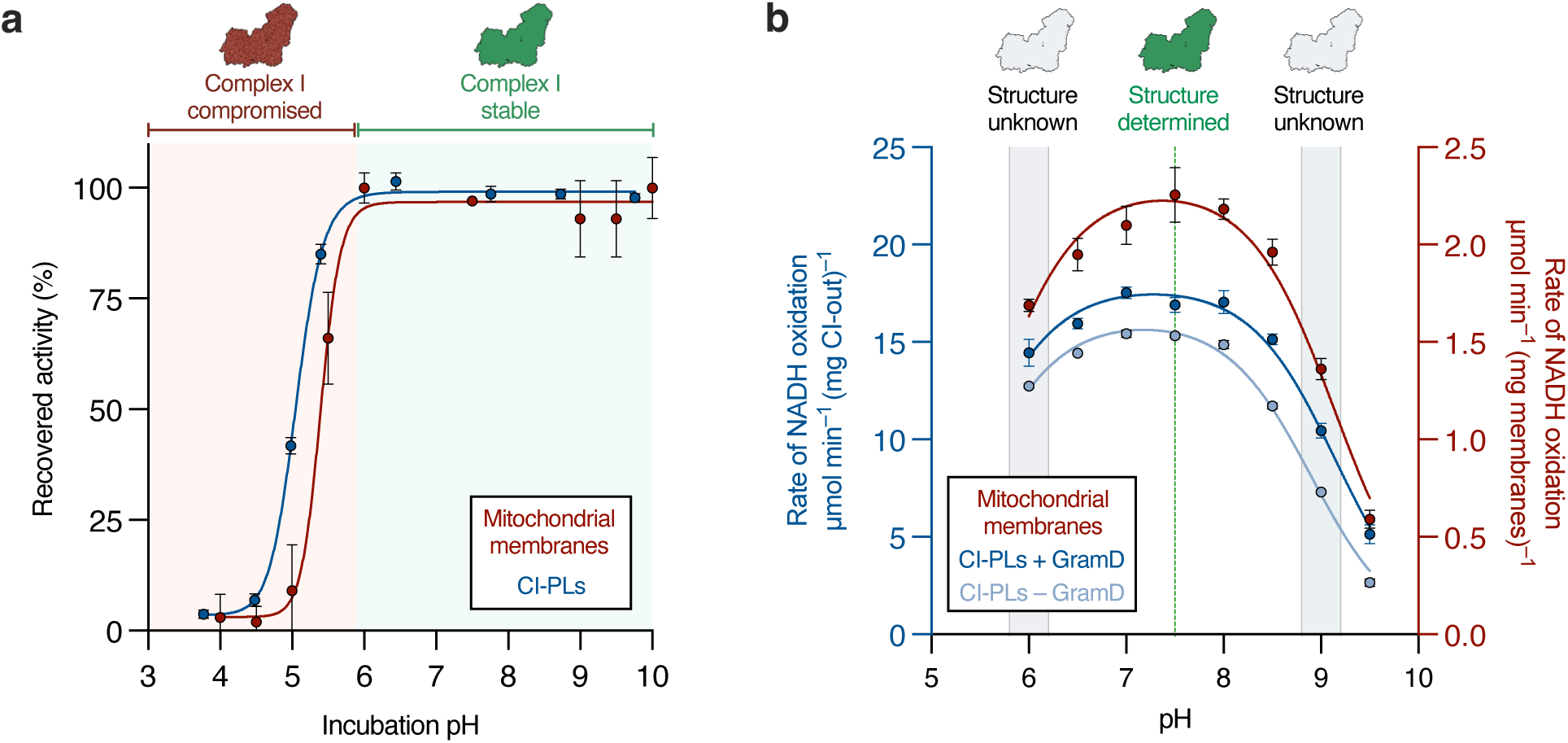
The pH dependence of complex I stability and catalysis. (**a**) The effect of pH on the stability of complex I from *B. taurus*. Mitochondrial membranes or complex I proteoliposomes (CI-PLs) were incubated for 1 hour on ice in CHB buffer (20 mM citric acid, 30 mM HEPES, 40 mM boric acid) at various pH values and the NADH:O_2_ oxidoreductase activity then determined at pH 7.5 in the presence of the alternative oxidase (AOX) from *Trypanosoma brucei brucei*^52^, relative to the sample with maximum activity. Complex I retained full activity at all pH values between 6.0 and 10.0, but was markedly unstable at lower pH values. (**b**) The NADH:O_2_ oxidoreductase activity of mitochondrial membranes and CI-PLs measured at different pH values in the presence of AOX^52^. Rates were measured in CHB buffer corrected to various pH values at the assay temperature (32 °C). Data for CI-PLs are shown with/without gramicidin D (GramD) to dissipate Δp. The bell-shaped curves can be described using two pKa values of 5.5 and 8.9. Structures of complex I from *B. taurus* have been solved at pH ∼7.5 (maximal rate of catalysis) but not at the upper and lower limits of stability and activity (pH ∼6.0 and ∼9.0). All data are mean averages with error (± S.D.) values from technical replicates (n = 3, mitochondrial membranes. n = 4, CI-PLs).

**Extended Data Fig. 2.**
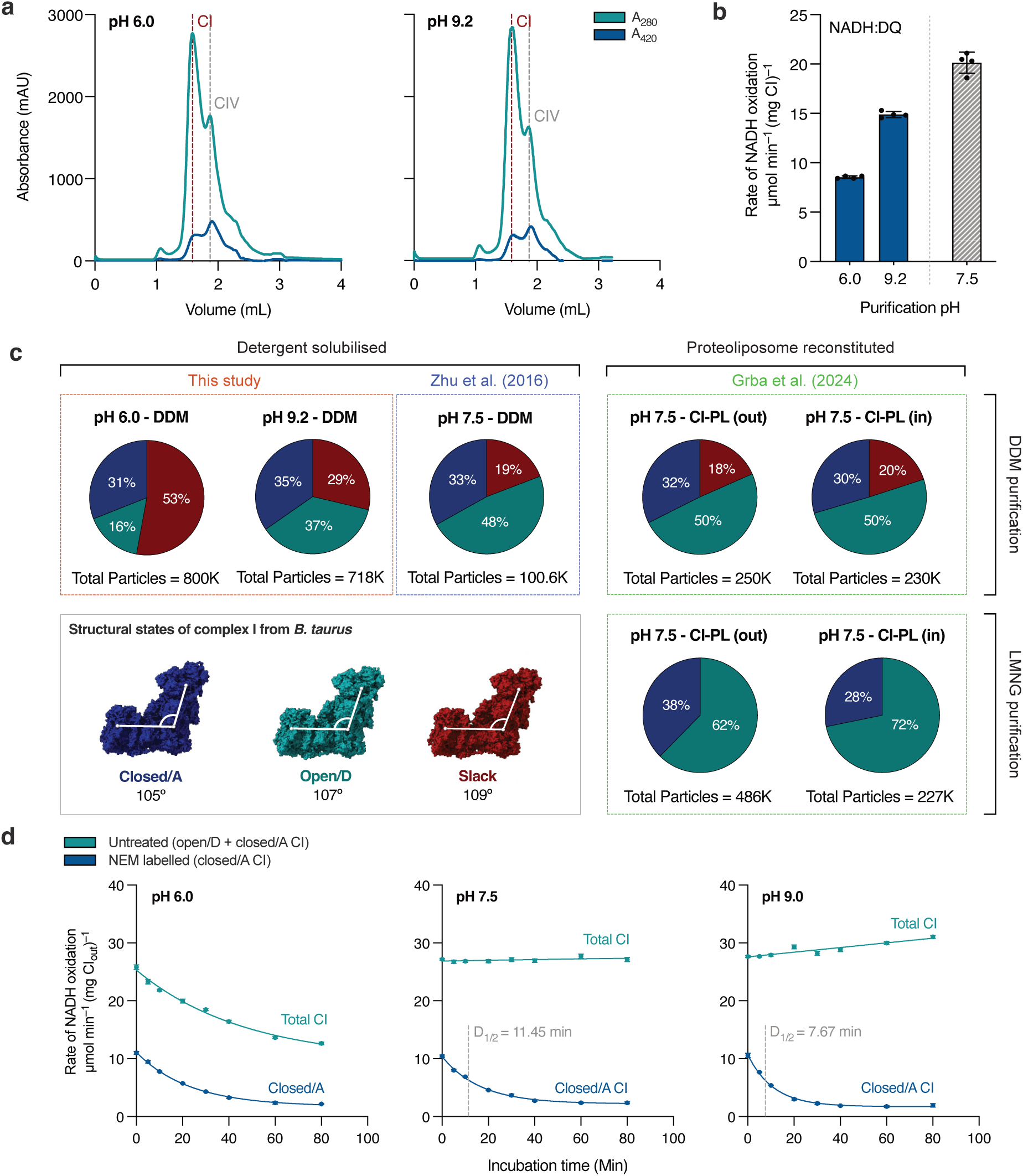
Biochemical characterization and global state distributions of complex I in cryo-EM samples. (**a**) Size exclusion chromatograms for cryo-EM samples at pH 6.0 and 9.2. Peaks corresponding to complex I and IV are indicated. (**b**) NADH:decylubiquinone oxidoreduction activities of the cryo-EM samples prepared at pH 6.0 and 9.2 (assays performed at pH 7.5). The activity values are shown relative to a typical complex I sample purified in DDM at pH 7.5 throughout (grey bar). Data are mean averages with error (± S.D.) values from technical replicates (n = 4) (**c**) Proportions of global resting states in cryo-EM analyses of bovine complex I. The proportion of closed/A (blue), open/D (green) and slack (red) states in complex I samples are shown for this study alongside a matching DDM-purified complex I sample at pH 7.5^9^, and proteoliposome reconstituted samples prepared at pH 7.5 from complex I purified either in DDM or LMNG^41^. Illustrative models of the three states of complex I from *B. taurus*, with the interdomain angle (measured between G334^ND5^, L111^ND1^ and FeS cluster N5) are given for reference. (**d**) Stability and deactivation of complex I in proteoliposomes incubated at pH 6.0, 7.5 and 9.0 at 30 °C. After the designated times of incubation, samples were moved onto ice, labelled with N-ethylmaleimide (NEM) and the NADH:O_2_ (+AOX) activity determined. The total activity is shown, alongside the activity after NEM treatment, which corresponds to only the closed/A enzyme present. Deactivation half-lives (D_1/2_) are given for pH 7.5 and 9.0. The enzyme is unstable at pH 6.0 at 30 °C. Data are mean averages with error (± S.D.) values from technical replicates (n = 3).

**Extended Data Fig. 3.**
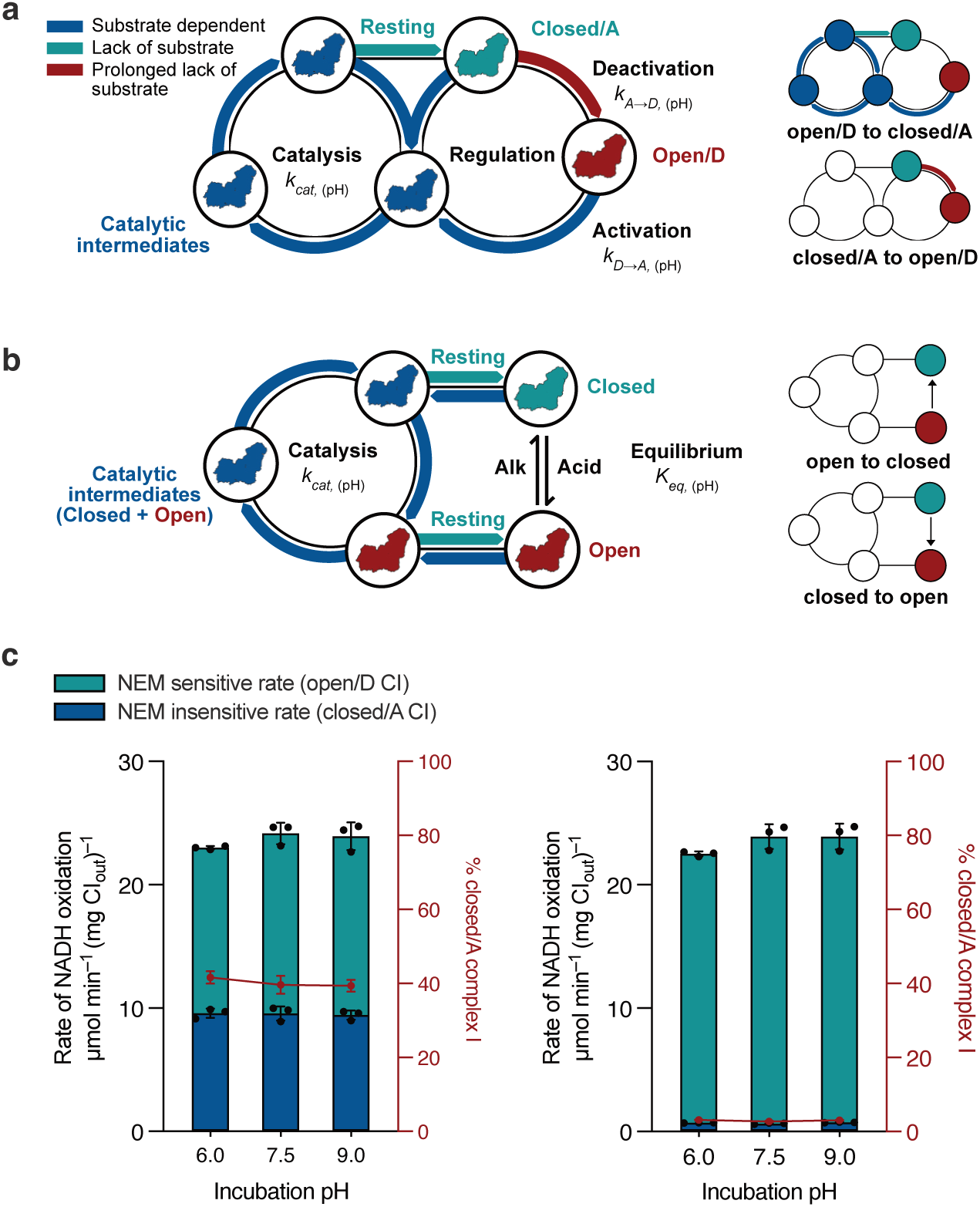
Resting state populations of bovine complex I are insensitive to pH^37–41^. (**a, b**) Schematic representations of conversions between the closed/A and open/D resting states. Mammalian complex I interconverts between two structurally distinct resting states: the ‘active’ structurally ‘closed’ state (closed/A), which is catalytically competent, and the ‘deactive’ structurally ‘open’ state (open/D), which requires substrate-induced reactivation^37–41^. When catalysis stops, for example in ischemic tissue, complex I first rests in closed/A. Then, during the ensuing warm ischemia (or thermal treatment *in vitro*) it relaxes into open/D. It returns to catalysis when substrates become available. **(a)** The widely accepted regulatory model in which activation/closure and deactivation/opening are unidirectional, condition-dependent processes and only closed states exist on the catalytic cycle^41^. **(b)** A recent alternative model^12^ in which a rapid, reversible, pH-dependent equilibrium exists between closed/A and open/D, even at low temperatures and without substrates, and both exist on the catalytic cycle. **(c)** The proportion of closed/A and open/D states determined by NEM sensitivity following incubation at pH 6.0, 7.5 and 9.0. The NEM assay uses the accessibility of C39^ND3^ to distinguish the states, as it is occluded in closed/A but accessible to NEM derivatization in open/D, where it prevents reactivation^45,71^. CI-PLs were diluted into CHB buffer (pH-corrected at 4 °C) and incubated on ice for 1 hour prior to activity measurements. For NEM treated samples, 1 mM NEM was included during incubation. NADH:O_2_ activity (+AOX) was then determined in proteoliposome buffer at pH 7.5. In the deactivated CI-PL samples, complex I was deactivated at pH 7.5 prior to diluting the CI-PLs into the CHB buffer. Note the essentially complete recovery of catalytic turnover in all cases. The proportion of closed/A did not respond to pH for either as-prepared or deactivated CI-PLs. As we showed previously that the NEM-determined proportions of closed/A in CI-PLs match exactly to the cryo-EM-determined particle populations^41^, we conclude that pH does not set the proportion of closed/A, supporting the regulatory scheme shown in panel a. Data are mean averages with error (± S.D.) values from technical replicates (n = 4).

**Extended Data Fig. 4.**
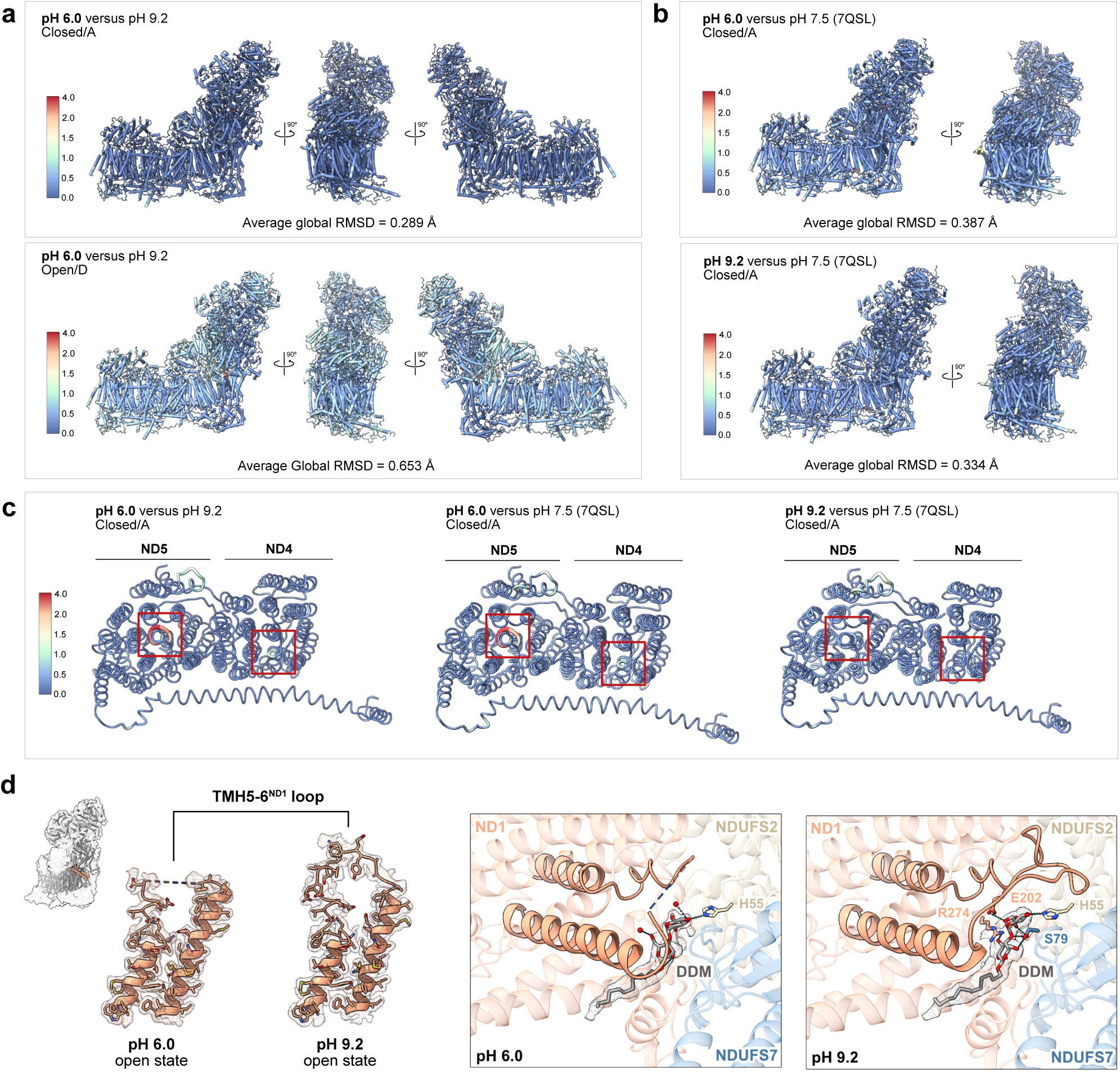
Comparison of complex I models at different pH values. (**a**) Alignment of closed/A states and open/D states at pH 6.0 and pH 9.2. Root-mean-square deviation (RMSD) values (Å) are given as global values. Bold font indicates the model shown in each case. (**b**) Comparison of the closed/A states from this study with the corresponding structures solved at pH 7.5 in nanodiscs (PDB: 7QSL). (**c**) Comparison of the antiporter-like subunits ND4 and ND5 at different pH values. Top-down views are shown with the colour scale and backbone width indicating the deviation between the models. The regions of highest variability between pH 6.0 and pH 9.2 structures, TMH8^ND5^ and TMH7^ND4^, are highlighted with red boxes. Variability in TMH7^ND4^ is associated with the alternate positions of the H220 sidechain. (**d**) The TMH5-6^ND1^ loop is disordered in open/D state complex I at pH 6.0 but adopts an ordered extended conformation at pH 9.2. Dashed lines indicate residues that are not modelled due to high flexibility and disordering. The position of TMH5-6^ND1^ is shown inset. DDM is bound close to the TMH5-6^ND1^ loop in open/D states at pH 6.0 and pH 9.2. Electron density for the DDM molecule is shown. Hydrogen bonding interactions between the DDM and local structural elements in ND1, NDUFS2 and NDUFS7 are indicated with dashed teal lines.

**Extended Data Fig. 5.**
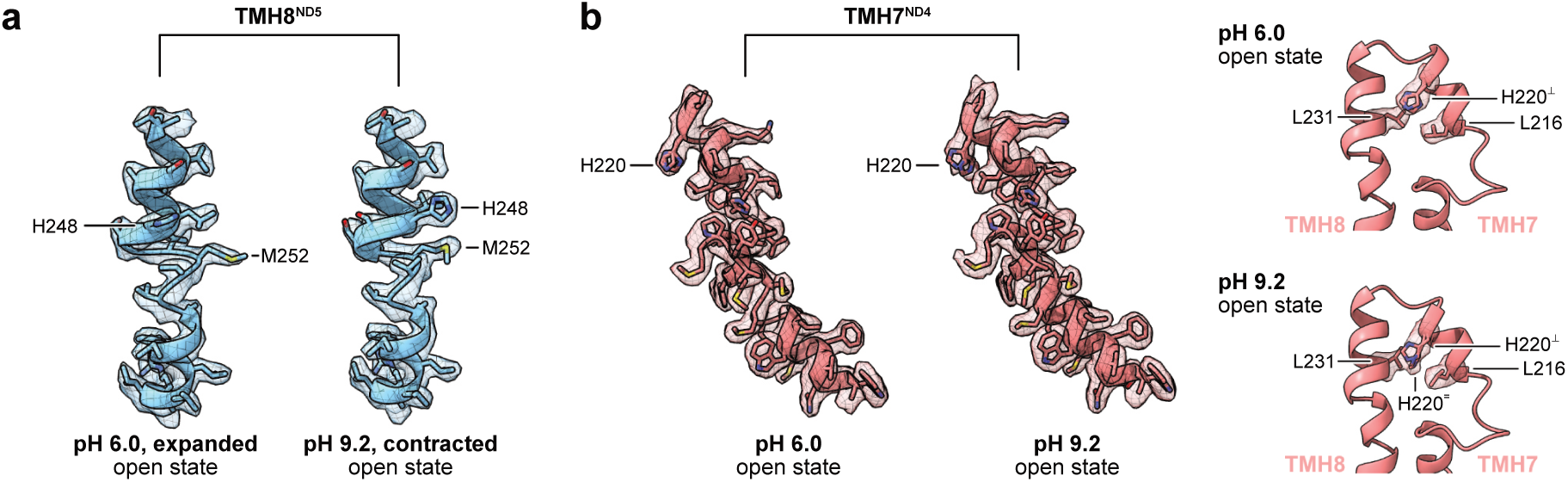
Conformational changes in ND5 and ND4 are conserved in the active/A and open/D states. (**a**) The pH dependent shift of the position of H248^ND5^ (see Fig. 1b for the same information for the closed/A state) is observed in the open/D state. (**b**) The electron density of TMH7^ND4^ in the open state/D (see Fig. 1c for the same information for the closed/A state). Only density corresponding to the perpendicular state of H220^ND4^ is observed in the pH 6.0 open/D state while density suggestive of both rotamers of H220^ND4^ is present at pH 9.2.

**Extended Data Fig. 6.**
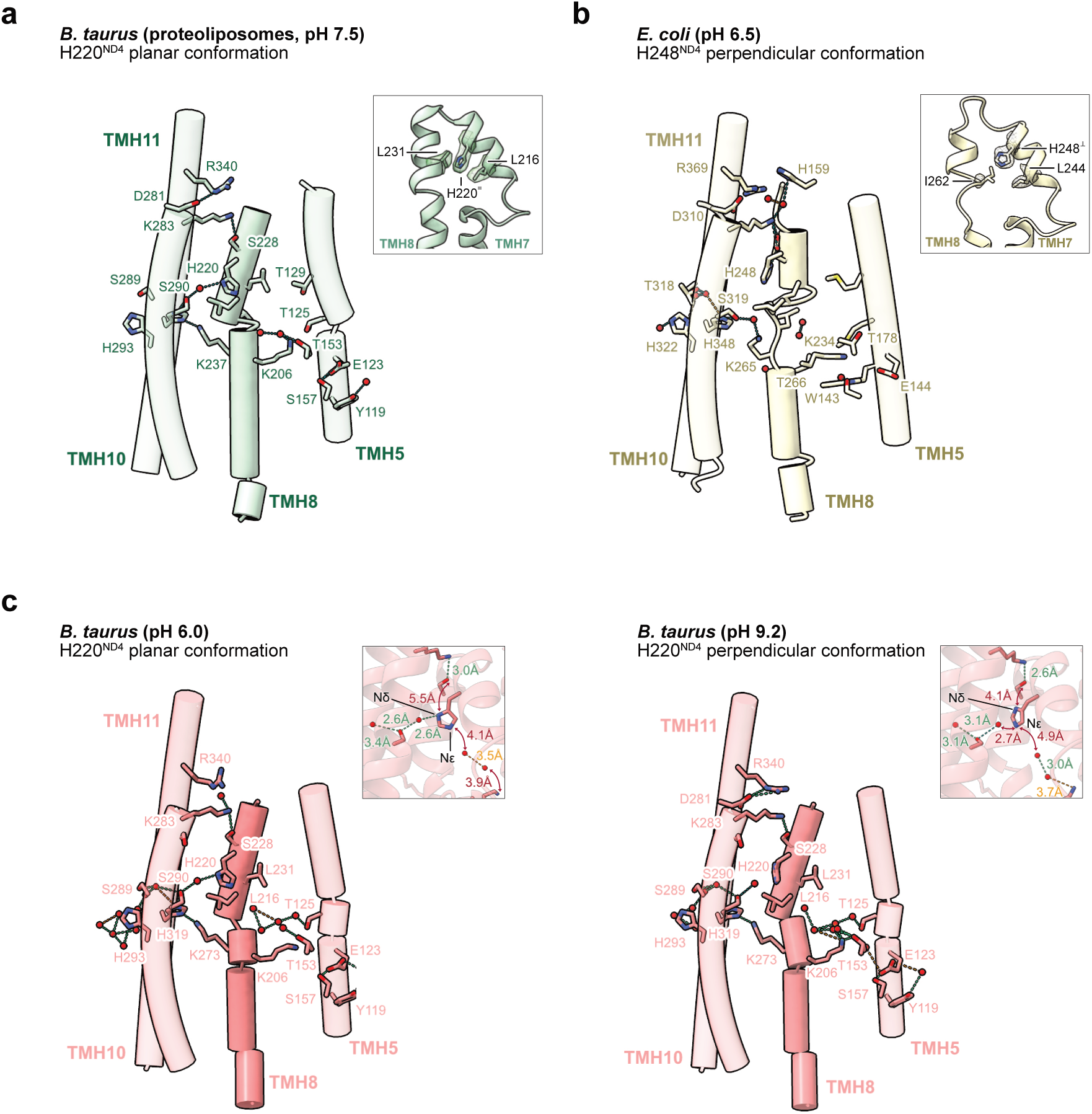
Analysis of proton transfer pathways in ND4 in additional structures. (a) Complex I from *B. taurus* proteoliposomes in the active state (PDB: 8Q48) shows the junction histidine (H220) solely in the planar conformation. Continuous connectivity between ND2 and ND5 is observed along the central axis. The electron density for H220 and the double leucine motif across TMH7 and TMH8 is shown inset, showing a single conformation could be modelled. (**b**) Complex I from *Escherichia coli* shows the junction histidine (H248) in a single perpendicular conformation. The electron density is shown inset for H248, with only one of the leucine residues conserved (L244) and the other replaced with an isoleucine (I262). (**c**) The alternative conformations of H220 in the structures from *B. taurus* at pH 6.0 (planar) and pH 9.2 (perpendicular). The conformations match those in Fig. 3 but slight modelling differences preclude a continuous hydrogen bonding network from ND2 to ND5 (planar conformation) or from the proton uptake channel to ND5 (perpendicular conformation). This is highlighted by the hydrogen bonding interactions of H220, shown inset, with distances that exceed 3.7 Å shown as red arrows.

**Extended Data Fig. 7.**
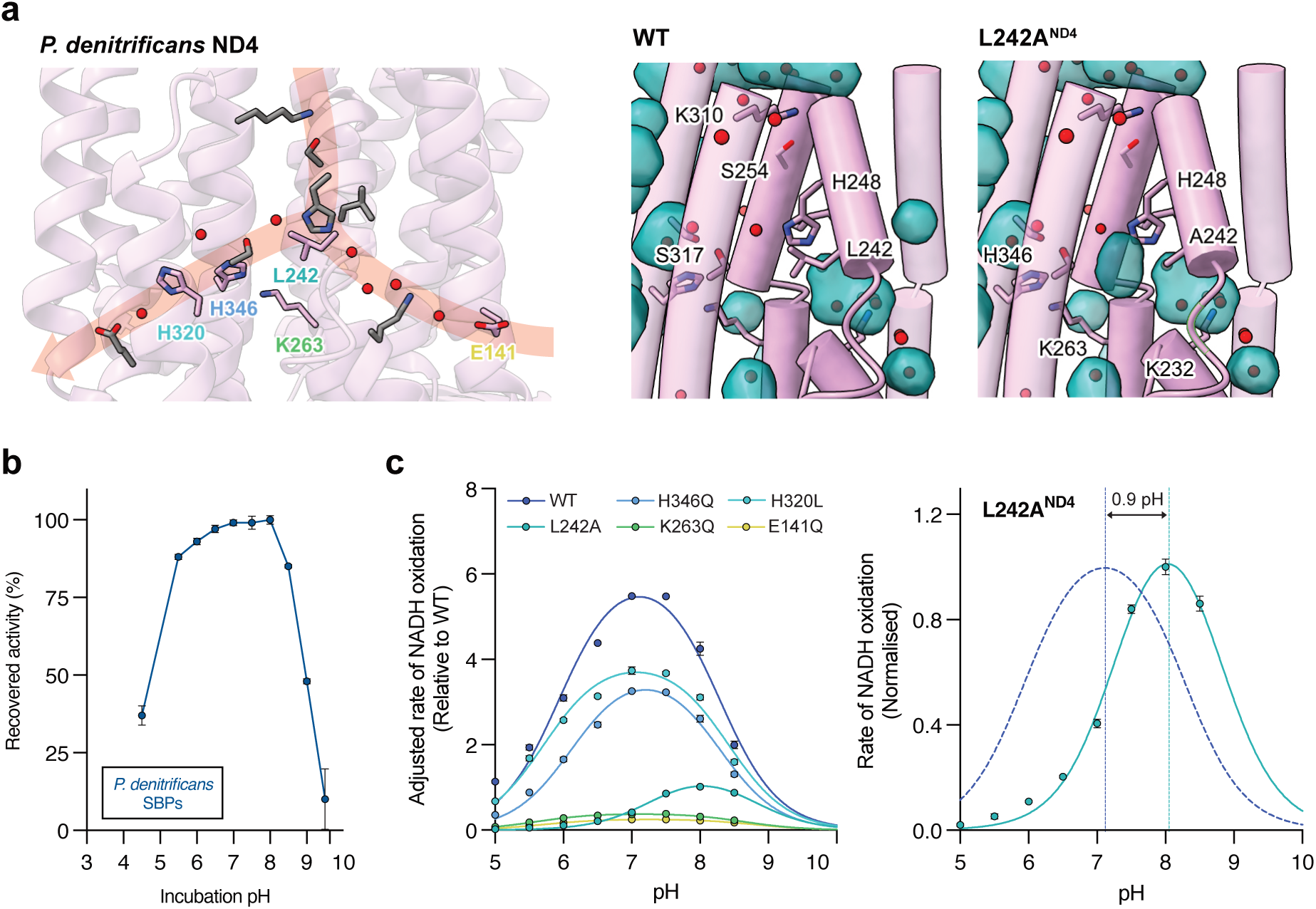
The pH dependence of *P. denitrificans* complex I activity in ND4 mutants. (**a**) Positions of mutated residues in the *P. denitrificans* ND4 subunit. Proposed proton transfer routes across the central axis and from the proton uptake channel are indicated. Predicted solvent access in the L242A mutant is shown on the right. Complex I from *P. denitrificans* was mutated in ChimeraX and cavities (green surfaces) were located using the built-in cavity finder. (**b**) Effect of pH on the stability of complex I-containing sub-bacterial membrane particles (SBPs) from *P. denitrificans*. The recovered activity (% of maximum) was determined after incubation in CHB buffer for 1 hour. (**c**) pH dependence of NADH:O_2_ oxidoreductase activity in ND4 mutants. Wild-type (WT) rates are determined in µmol NADH min^−1^ (mg SBP)^−1^. Rates of mutants were adjusted to complex I content as determined by NADH:APAD^+^ oxidoreductase activity at the flavin site for NADH oxidation, and displayed relative to the WT. SBPs were treated with 0.5 µg AOX per µg SBPs (see Methods) and uncoupled with 1 µg mL^−1^ gramicidin D. The L242A mutant shifts the optimum pH for catalysis by 0.9 pH units. L242A activity is shown normalised against the WT activity (dashed blue line) to highlight the change in optimum pH (right). Data are mean averages with error (± S.D.) values from technical replicates (n = 3).

**Extended Data Fig. 8.**
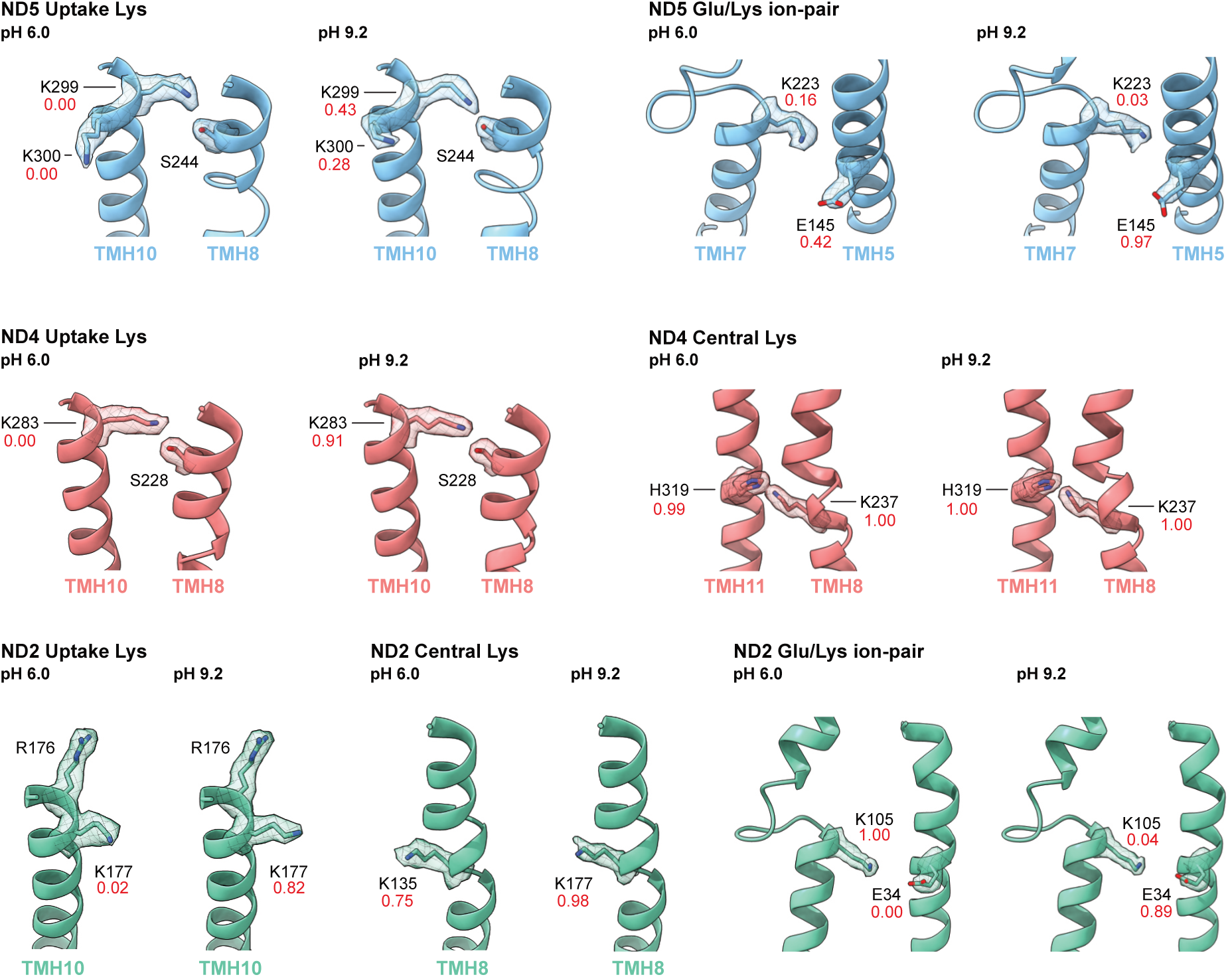
Cryo-EM densities for key residues predicted in CpHMD simulations to change proton state between pH 6.0 and G.2. The predicted proportions of deprotonation (from Fig. 4) are given for reference.

**Extended Data Fig. 9.**
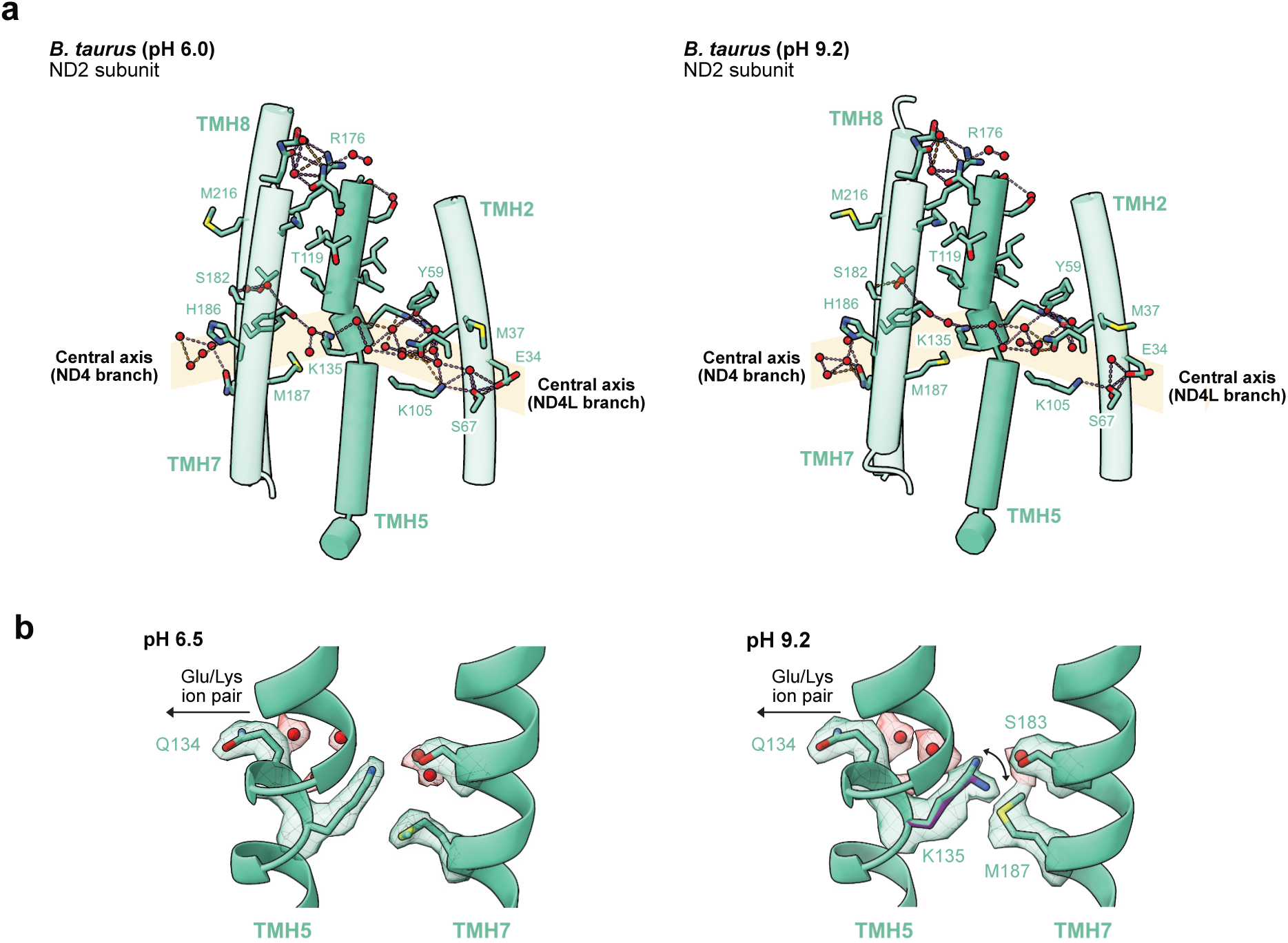
Analysis of proton transfer connectivity in the ND2 subunit. (**a**) ND2 in pH 6.0 and pH 9.2 shows connectivity across the subunit from ND4L to ND4. For ease of comparison, transmembrane helices are labelled according to their analogous helices in ND4 and ND5. Only minor differences are observed between pH values in the water molecules resolved and the positions of the sidechains for M37, M187 and M216, which are not expected to be directly involved in proton transfer. (**b**) Possible pH dependent conformational changes around the central lysine (K135) in ND2. The pH 6.0 structure suggest a single position of the K135 sidechain which is positioned to interact with the water network (red spheres, with red density) that connects to the Glu/Lys ion pair at the entrance of the subunit. A Y-shaped feature in the pH 9.2 density map suggests a variation in sidechain orientation such that a second rotamer can be modelled into the density, pointing away from the water network to the Glu/Lys ion pair and no longer forming a continuous network across the subunit. The sidechain of M187 also rotates at this pH.

