## Supplementary Information for "Conformational gating at histidine junctions coordinates proton translocation in respiratory complex I"

**This file includes:**
Supplementary Tables 1-5
Supplementary Figures 1-7

**Supplementary Table 1: Cryo-EM refinement and validation statistics for the pH 6.0 dataset**

| | **Closed/A state** | **Open/D state** |
|---|---|---|
| **Map statistics** | | |
| Final sampling rate (Å/px) | 0.798 | 0.798 |
| Symmetry imposed | *C*1 | *C*1 |
| Final particle images | 247,586 | 128,687 |
| Map resolution (Å) (FSC) | 2.258 (0.143) | 2.494 (0.143) |
| Map resolution range (Å) | 1.945 – 4.723 | 2.162 – 5.025 |
| Map sharpening B-factor (Å$^2$) | -23.8 | -18.8 |
| **Model statistics** | | |
| Initial model used | 7QSL | 7QSN |
| Model resolution (Å) (FSC) | 2.3 (0.5) | 2.6 (0.5) |
| *Model composition* | | |
| Nonhydrogen atoms | 70,176 | 70,260 |
| Protein residues | 8,270 | 8,222 |
| Ligands | 57 | 61 |
| Waters | 1,637 | 1,879 |
| *Mean B-factors (Å$^2$)* | | |
| Protein | 48.52 | 67.30 |
| Ligand | 65.44 | 93.01 |
| Water | 43.02 | 67.16 |
| *RMS deviations* | | |
| Bond lengths (Å) | 0.008 | 0.011 |
| Bond angles (°) | 0.708 | 0.749 |
| **Validation statistics** | | |
| MolProbity score | 1.87 | 2.06 |
| Clashscore | 5.83 | 8.59 |
| Rotamer outliers (%) | 2.51 | 1.93 |
| Cß outliers (%) | 0.00 | 0.00 |
| *Ramachandran plot* | | |
| Favoured (%) | 96.32 | 94.48 |
| Allowed (%) | 3.42 | 5.06 |
| Outliers (%) | 0.26 | 0.46 |
| *Rama-Z score (RMSD)* | | |
| Whole | -2.63 (0.08) | -3.18 (0.08) |
| Helix | -2.20 (0.06) | -2.48 (0.06) |
| Sheet | -0.71 (0.23) | -0.95 (0.25) |
| Loop | -0.96 (0.10) | -1.49 (0.10) |

**Supplementary Table 2: Cryo-EM refinement and validation statistics for the pH 9.2 dataset**

| | **Closed/A state** | **Open/D state** |
|---|---|---|
| **Map statistics** | | |
| Final sampling rate (Å/px) | 0.800 | 0.800 |
| Symmetry imposed | *C*1 | *C*1 |
| Final particle images | 249,146 | 262,659 |
| Map resolution (Å) (FSC) | 2.243 (0.143) | 2.308 (0.143) |
| Map resolution range (Å) | 1.903 – 4.673 | 1.988 – 5.003 |
| Map sharpening B-factor (Å$^2$) | -21.1 | -21.2 |
| **Model statistics** | | |
| Initial model used | 7QSL | 7QSN |
| Model resolution (Å) (FSC) | 2.3 (0.5) | 2.4 (0.5) |
| *Model composition* | | |
| Nonhydrogen atoms | 70,419 | 70,327 |
| Protein residues | 8,270 | 8,224 |
| Ligands | 57 | 60 |
| Waters | 1,834 | 1,916 |
| *Mean B-factors (Å$^2$)* | | |
| Protein | 42.46 | 62.42 |
| Ligand | 60.16 | 84.99 |
| Water | 39.84 | 64.42 |
| *RMS deviations* | | |
| Bond lengths (Å) | 0.008 | 0.002 |
| Bond angles (°) | 0.698 | 0.513 |
| **Validation statistics** | | |
| MolProbity score | 1.82 | 1.55 |
| Clashscore | 5.33 | 5.01 |
| Rotamer outliers (%) | 2.76 | 1.11 |
| Cß outliers (%) | 0.00 | 0.00 |
| *Ramachandran plot* | | |
| Favoured (%) | 96.78 | 96.24 |
| Allowed (%) | 3.07 | 3.48 |
| Outliers (%) | 0.16 | 0.28 |
| *Rama-Z score (RMSD)* | | |
| Whole | -2.27 (0.08) | -2.81 (0.08) |
| Helix | -1.89 (0.06) | -2.33 (0.05) |
| Sheet | -0.55 (0.24) | -0.97 (0.24) |
| Loop | -0.81 (0.10) | -1.04 (0.10) |

**Supplementary Table 3: Comparison of the junction histidine positions in subunits ND4 and ND5 in structures of complex I from different species.** Representative structures were selected on the basis of resolution and map quality, with globally closed structures analysed where possible. Multiple instances of bovine complex I were included for comparison to the structures determined here. The position of $H220^{ND4}$ is only considered for resolutions < 2.5 Å due to the subtle nature of the variation. The conformation of $TMH8^{ND5}$ can be distinguished at lower resolutions and is considered in all cases. The positions given are those modelled in the original datasets, with instances where alternative conformations could be modelled based on the density noted.

| Species | State | | PDB | Resolution (Å) | pH | Preparation | ND5 His position | ND4 His position |
|---|---|---|---|---|---|---|---|---|
| | Given name | Global | | | | | | |
| **Mammals** | | | | | | | | |
| *Bos taurus* | pH 6 active | Closed | 9R12 | 2.2 | 6.0 | Detergent (DDM) | Expanded (H248) | Perpendicular/dual occupancy (H220) |
| | pH 9 active | Closed | 9R14 | 2.2 | 9.2 | Detergent (DDM) | Contracted (H248) | Dual occupancy (H220) |
| | pH 6 deactive | Open | 9R13 | 2.5 | 6.0 | Detergent (DDM) | Expanded (H248) | Perpendicular (H220) |
| | pH 9 deactive | Open | 9R15 | 2.3 | 9.2 | Detergent (DDM) | Contracted (H248) | Dual Occupancy (H220) |
| | Active-Q10 | Closed | 7QSK | 2.8 | 7.5 | Nanodisc (from DDM) | Contracted (H248) | --- |
| | Deactive (composite) | Open | 7QSM | 2.3 | 7.5 | Nanodisc (from DDM) | Contracted (H248) | Planar[1] (H220) |
| | IM1761092 active | Closed | 7R41 | 2.3 | 7.1 | Detergent (DDM) | Expanded (H248) | Perpendicular[1] (H220) |
| | IM1761092 deactive | Open | 7R45 | 2.4 | 7.1 | Detergent (DDM) | Expanded (H248) | Perpendicular (H220) |
| | CI-out, active | Closed | 8Q48 | 2.5 | 7.5 | Proteoliposomes (from LMNG) | Contracted (H248) | Planar (H220) |
| *Mus musculus* | Active | Closed | 8OM1 | 2.5 | 7.2 | Detergent (DDM) | Contracted (H248) | Perpendicular (H220) |
| *Ovis aries* | Turnover-closed | Closed | 6ZKC | 3.1 | 7.4 | Detergent (LMNG) | Contracted (H248) | --- |
| *Sus scrofa* | Active-Q10 | Closed | 7V2C | 2.9 | 7.5 | Detergent (digitonin) | Contracted (H248) | --- |
| | In situ, composite | Closed | 8UD1 | 2.1 | 7.8 | In situ | Contracted (H248) | Planar[1] (H220) |
| **Insect** | | | | | | | | |
| *Drosophila melanogaster* | Active (Dm1) | Closed | 8B9Z | 3.3 | 7.8 | Detergent (DDM) | Intermediate (H221) | --- |
| | Helix-locked | Closed | 8ESZ | 3.4 | 7.8 | Amphipol (from digitonin) | Expanded (H221) | --- |
| **Bacteria** | | | | | | | | |
| *Paracoccus denitrificans* | Active | Closed | 8QBY | 2.3 | 6.5 | Nanodisc (from DDM) | Expanded (H263) | Planar (H246) |
| *Escherichia coli* | Turnover-closed | Closed | 7Z7S | 2.4 | 6 | Detergent (LMNG) | Expanded (H254) | Perpendicular (H248) |
| *Mycobacterium smegmatis* | Both Q modelled | Closed | 8E9G | 2.6 | 6.0 | Detergent (DDM) | Expanded (H256) | Not conserved |
| *Thermus thermophilus* | Entire complex | Open | 4HEA | 3.3 | 6.0 | Crystal (TDM) | Expanded (H241) | --- |
| **Yeasts** | | | | | | | | |
| *Yarrowia lipolytica* | Deactive | Open | 6YJ4 | 2.7 | 7.4 | Detergent (DDM) | Contracted (H251) | --- |
| *Chaetomium thermophilum* | State 2 | Closed | 7ZMB | 2.8 | 7.4 | Detergent (LMNG) | Contracted (H248) | --- |
| *Pichia pastoris* | State 1 | Closed | N/A | 2.8 | 7.4 | Nanodisc (from LMNG) | Expanded (H248) | --- |
| **Protists** | | | | | | | | |
| *Tetrahymena thermophila* | CI+CIII SC | Closed | 7TGH | 2.6 | 7.4 | Amphipol (from digitonin) | Expanded (H430) | --- |
| *Euglena gracilis* | Turnover | Closed | 8J9I | 2.9 | 7.4 | Detergent (LMNG) | Expanded (H257)[1] | --- |
| *Polytomella sp.* | Closed | Closed | 7ARD | 3.1 | 7.4 | Detergent (LMNG) | Expanded (H243) | --- |
| **Plants** | | | | | | | | |

| | | | | | | | | |
|---|---|---|---|---|---|---|---|---|
| *Arabidopsis thaliana* | CI-CIII SC | Closed | 8BPX | 2 | 7.4 | Detergent (GDN) | Contracted (H257) | Planar[1] (H241) |
| *Vigna radiata* | CI-CIII SC | Intermediate | 8E73 | 3.2 | 7.7 | Amphipol (from digitonin) | Contracted (H257) | --- |
| **Other/homologues** | | | | | | | | |
| *Bacillus pseudofirmus* | Mrp antiporter | N/A | 7QRU | 2.2 | 8.0 | Detergent (LMNG) | Dual conformation | N/A |

[1]An alternative conformation could also be modelled into the density.

**Supplementary Table 4:** Composition, total time and treatment of titratable residues in MD simulations.

<table>
<tr><th>Simulation type</th><th>Model</th><th>Inner water</th><th>Ext. water</th><th>$Na^+$</th><th>$Cl^-$</th><th>Buffer</th><th>Time (ns)</th><th>Titratable Residues</th></tr>
<tr><td rowspan="2">Initial equilibration</td><td>pH 6</td><td>315</td><td>22695</td><td>62</td><td>37</td><td>0</td><td>161</td><td rowspan="2">Adjusted to neutral pH</td></tr>
<tr><td>pH 9.2</td><td>364</td><td>22669</td><td>62</td><td>37</td><td>0</td><td>156</td></tr>
<tr><td rowspan="2">CpHMD</td><td>pH 6</td><td>315</td><td>22695</td><td>33</td><td>23</td><td>42</td><td>340+252+258</td><td rowspan="2">Titrated</td></tr>
<tr><td>pH 9.2</td><td>364</td><td>22669</td><td>33</td><td>23</td><td>42</td><td>334+254+240</td></tr>
<tr><td rowspan="2">Metadynamics</td><td>pH 6</td><td>315</td><td>22695</td><td>47</td><td>23</td><td>0</td><td>264 (HSE)<br>390 (HSD)</td><td rowspan="2">Fixed to CpHMD values</td></tr>
<tr><td>pH 9.2</td><td>364</td><td>22669</td><td>53</td><td>23</td><td>0</td><td>268 (HSE)<br>381 (HSD)</td></tr>
</table>

**Supplementary Table 5:** Titrated residues in CpHMD simulations and their respective λ-bias barrier (kJ mol$^{-1}$). The protonation states fixed in the metadynamics simulations for both pH models are denoted as 0 for protonated and 1 for deprotonated, and HSD/HSE shows the His tautomer used.

| CpHMD | | | Metadynamics | |
|---|---|---|---|---|
| Subunit | Residue | Barrier | pH 6.0 | pH 9.2 |
| ND5 | E145 | 10 | 0 | 1 |
| ND5 | D179 | 7.5 | 1 | 1 |
| ND5 | K223 | 12.5 | 0 | 0 |
| ND5 | H248 | 10 | HSD | HSD |
| ND5 | K299 | 7.5 | 0 | 1 |
| ND5 | K300 | 7.5 | 0 | 0 |
| ND5 | H328 | 7.5 | HSE | HSE |
| ND5 | H332 | 7.5 | HSE | HSE |
| ND5 | K336 | 7.5 | 1 | 1 |
| ND5 | K392 | 7.5 | 0 | 0 |
| ND5 | D393 | 7.5 | 1 | 1 |
| ND5 | E397 | 7.5 | 1 | 1 |
| ND5 | D554 | 7.5 | 1 | 1 |
| ND5 | E559 | 7.5 | 1 | 1 |
| ND5 | K564 | 7.5 | 0 | 0 |
| ND5 | H605 | 7.5 | HSE | HSE |
| ND5 | E606 | 7.5 | 0 | 1 |
| ND4 | E123 | 15 | 1 | 1 |
| ND4 | E141 | 7.5 | 1 | 1 |
| ND4 | K206 | 10 | 0 | 0 |
| ND4 | H213 | 7.5 | HSE | HSE |
| ND4 | K218 | 7.5 | 0 | 0 |
| ND4 | H220 | 10 | HSD/HSE | HSD/HSE |
| ND4 | E222 | 7.5 | 1 | 1 |
| ND4 | K237 | 10 | 1 | 1 |
| ND4 | K283 | 7.5 | 0 | 1 |
| ND4 | H293 | 10 | HSE | HSE |
| ND4 | H319 | 7.5 | HSE | HSE |
| ND4 | E378 | 7.5 | 1 | 1 |

| CpHMD | | | Metadynamics | |
|---|---|---|---|---|
| Subunit | Residue | Barrier | pH 6.0 | pH 9.2 |
| ND2 | E34 | 7.5 | 0 | 1 |
| ND2 | E54 | 7.5 | 1 | 1 |
| ND2 | K58 | 7.5 | 0 | 0 |
| ND2 | K105 | 10 | 1 | 0 |
| ND2 | E117 | 7.5 | 1 | 1 |
| ND2 | K135 | 10 | 0 | 1 |
| ND2 | K177 | 7.5 | 0 | 1 |
| ND2 | H186 | 7.5 | HSE | HSE |
| ND2 | K263 | 10 | 0 | 0 |
| ND2 | E269 | 7.5 | 1 | 1 |
| ND2 | K272 | 7.5 | 0 | 0 |
| ND2 | E347 | 7.5 | 0 | 1 |
| ND4L | E70 | 10 | 0 | 0 |

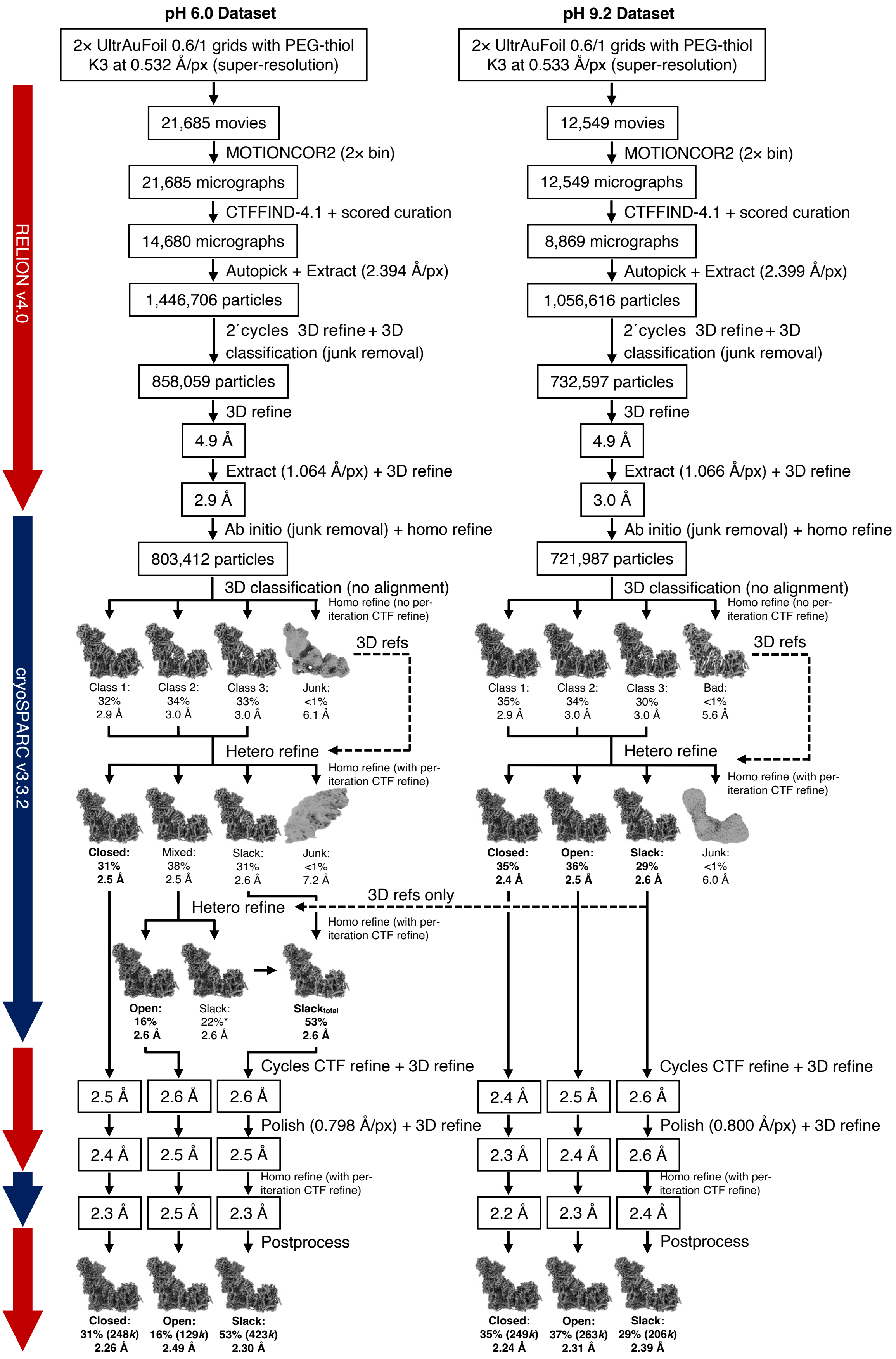


**Supplementary Fig. 1. High-resolution image-processing pipeline for cryo-EM datasets of bovine complex I at pH 6.0 and 9.2.** All early junk-particle removal using RELION was for classes with no morphological resemblance to complex I, and all major (>1%) complex I-like classes were included in classifications. It was necessary to further separate the pH 6 open/D and slack states using heterogeneous refinement.

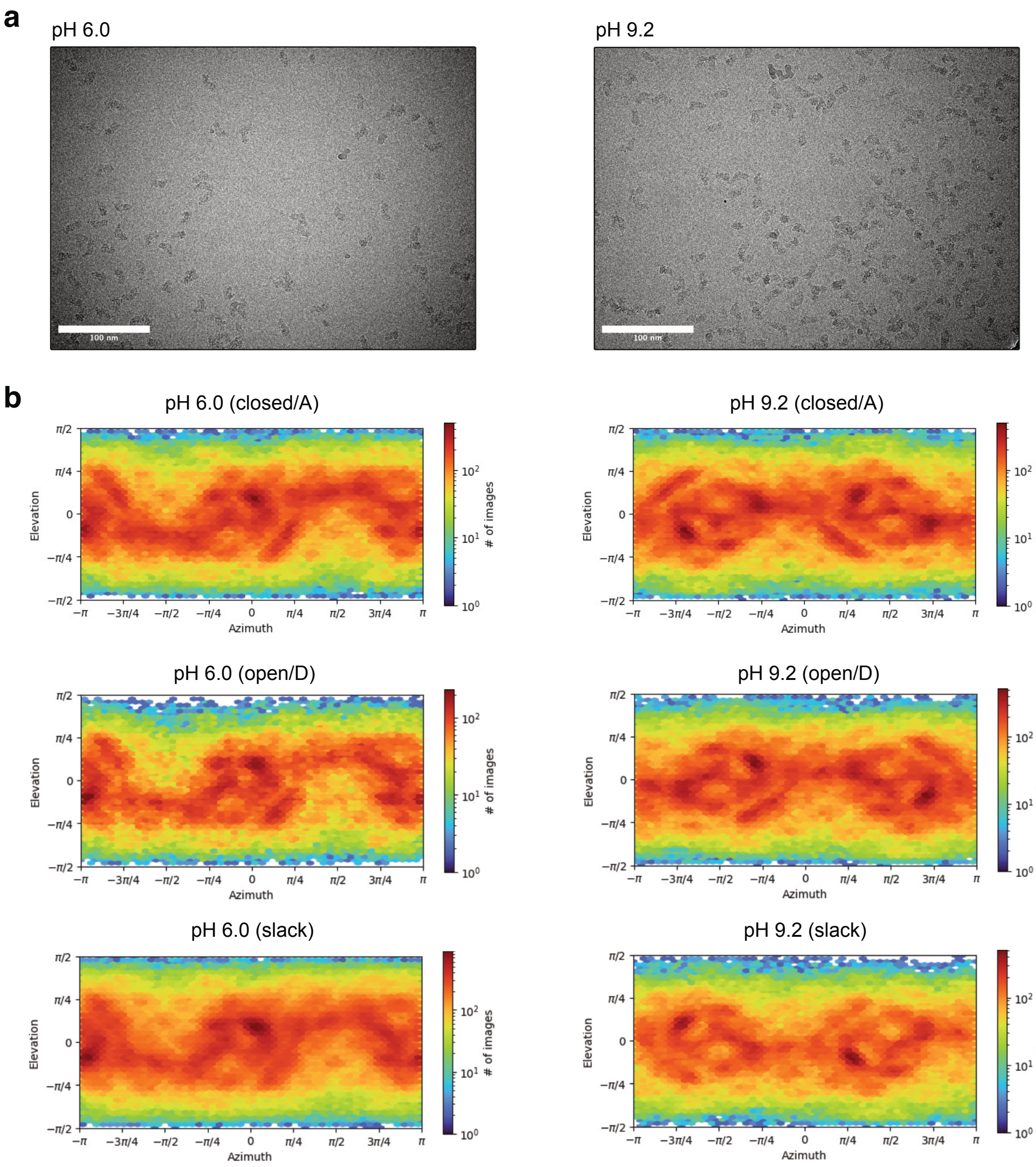


**Supplementary Fig. 2. Representative micrographs and cryo-EM particle orientation distributions for the particles used to construct the global maps.** (**a**) Representative cryoEM micrographs from the pH 6.0 and pH 9.2 datasets. (**b**) Azimuth-elevation plots of particle views are shown for each complex I state. Plots were generated in cryoSPARC v3.3.2.

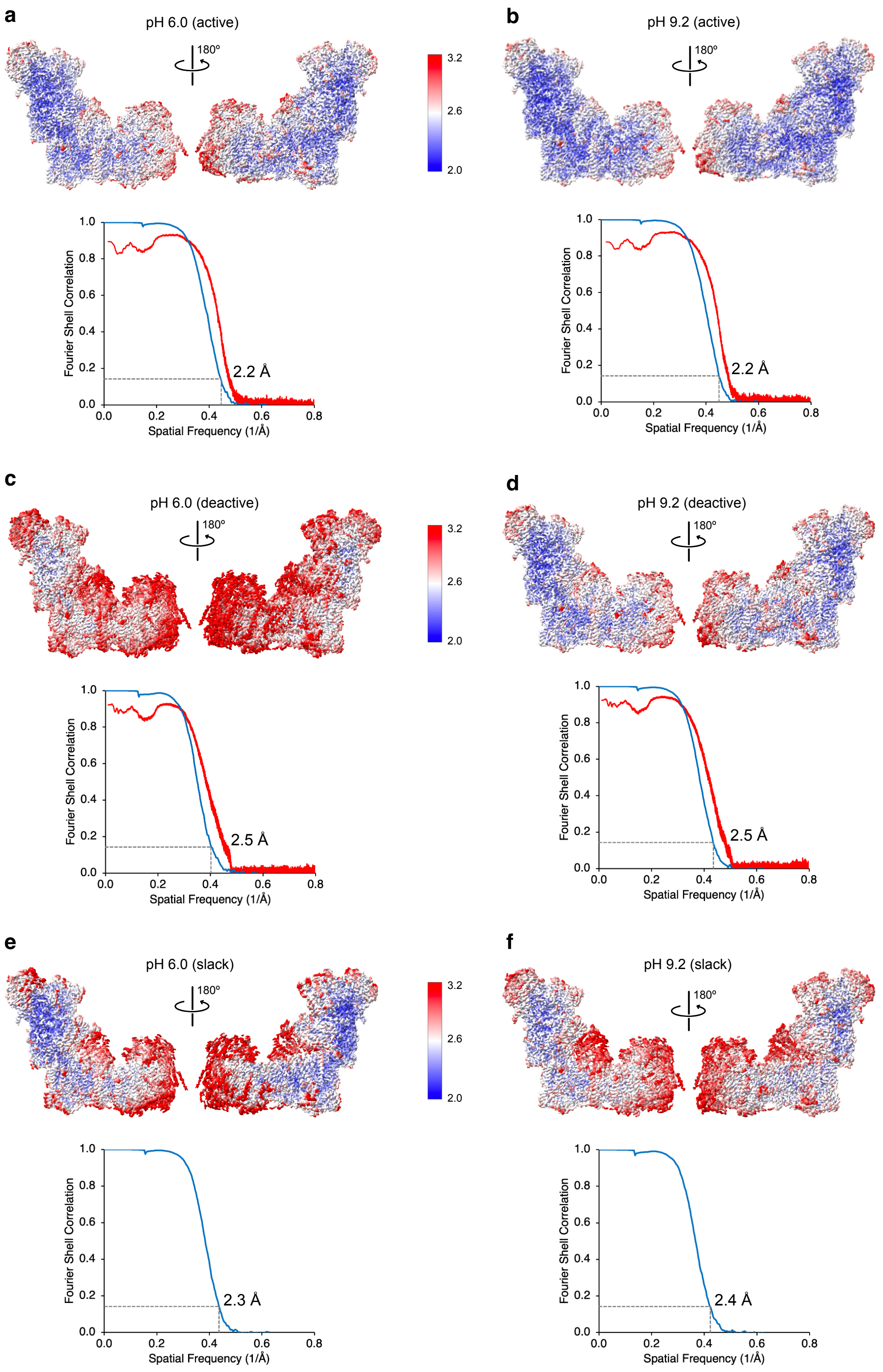


**Supplementary Fig. 3. Global and local resolution estimates for global maps and models.** Local resolution maps (FSC cutoff = 0.5), FSC curves (from two independent half-maps with FSC cutoff = 0.143, blue) and masked map-to-model FSC curves (where relevant, red) are shown for: (**a**) pH 6.0 closed/A state; (**b**) pH 9.2 closed/A state; (**c**) pH 6.0 open/D state; (**d**) pH 9.2 open/D state; (**e**) pH 6.0 slack state; (**f**) pH 9.2 slack state. Local resolutions were estimated using CryoSPARC v3.3.2 and visualized using UCSF ChimeraX 1.8. FSC curves were generated using RELION-4.0 postprocessing. Map-to-model FSC curves were generated using Mtriage in PHENIX v1.21.

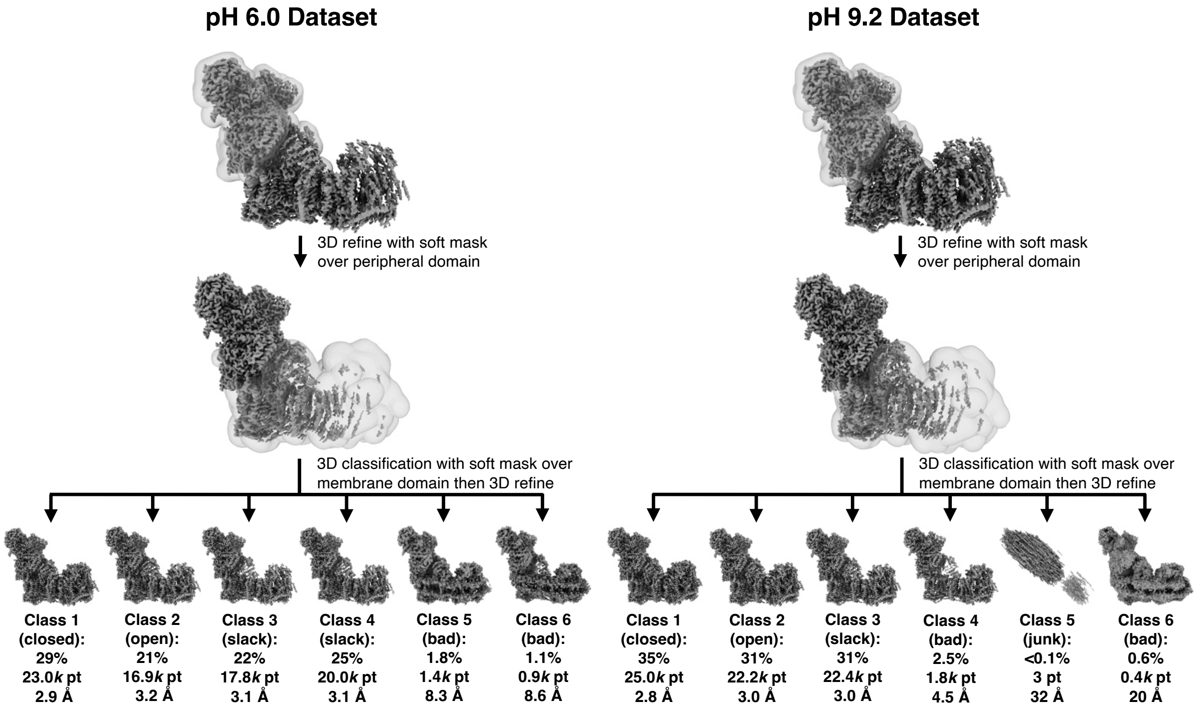


**Supplementary Fig. 4. Focus-Revert-Classify approach to validate global classification schemes.** Focus-Revert-Classify on a 10% particle subset on RELION v4.0 yielded one closed/A state for each dataset, plus three (pH 6.0) and two (pH 9.2) further states. One of these states from each dataset was identified as the open/D state, and the remaining two (pH 6.0) and one (pH 9.2) states were identified as the 'slack' state. The remaining low-population low-resolution classes were dismissed. The proportions of closed/A state particles were within 2% of the values obtained from the global classification scheme.

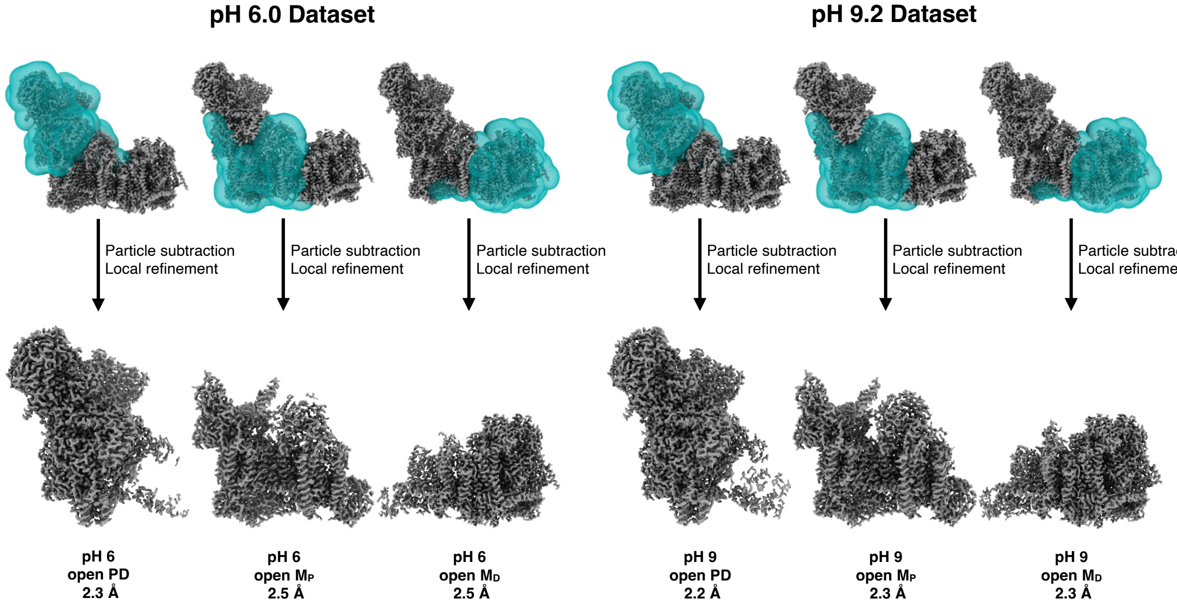


**Supplementary Fig. 5. Generation of focused maps for the pH 6.0 and 9.2 open/D states.** Focused maps for the peripheral domain (PD), proximal membrane domain ($M_P$) and distal membrane domain ($M_D$) of the open/D states at pH 6.0 (left) and pH 9.2 (right) were generated using soft masks and then combined to generate composite maps for modelling.

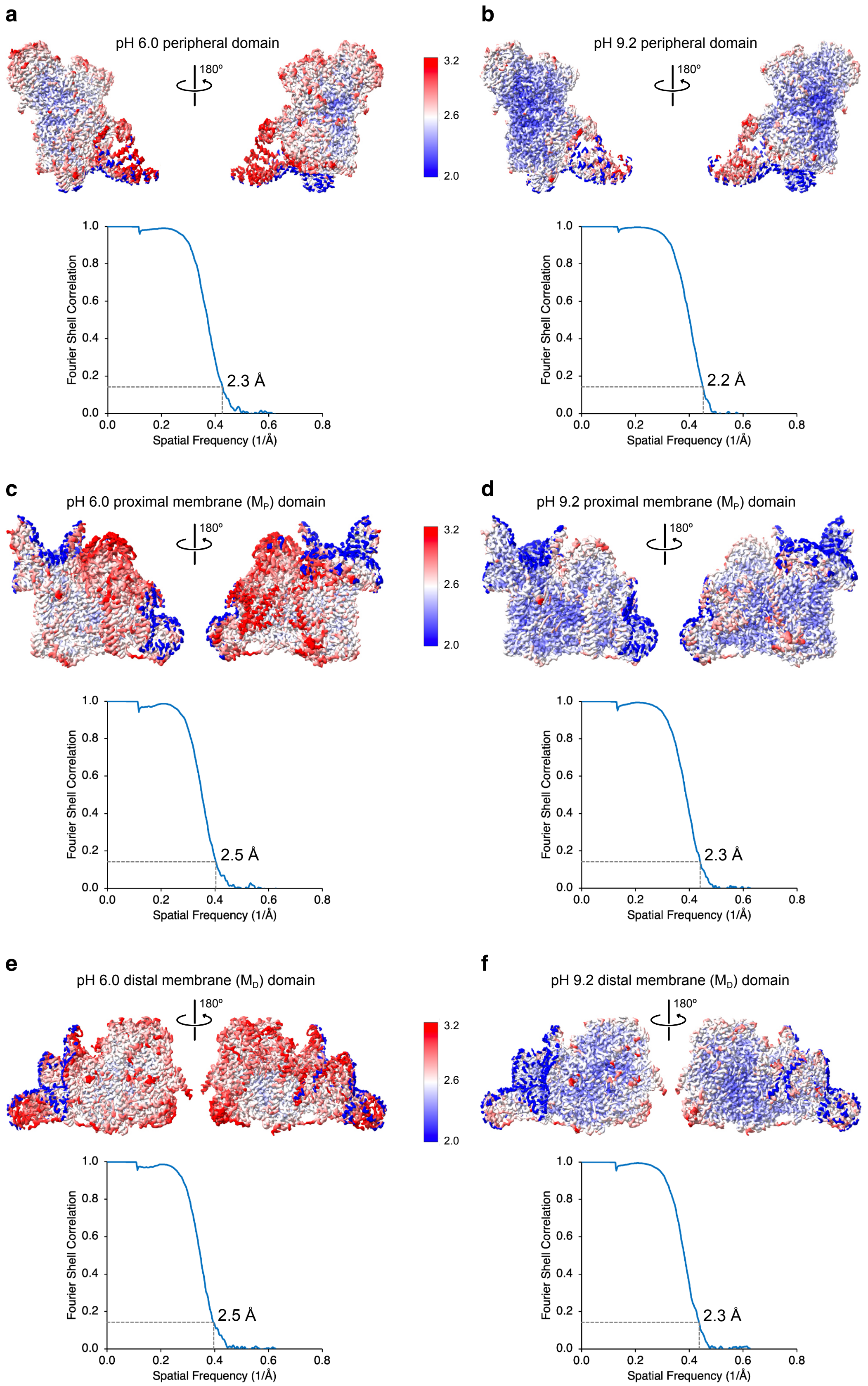


**Supplementary Fig. 6. Global and local resolution estimates for the focused maps used to generate the composite maps for the open/D state.** Local resolution maps (FSC cutoff = 0.5) and FSC curves (from two independent half-maps with FSC cutoff = 0.143, blue) are shown for: (**a**) pH 6.0 peripheral domain; (**b**) pH 9.2 peripheral domain; (**c**) pH 6.0 proximal membrane ($M_P$) domain; (**d**) pH 9.2 $M_P$ domain; (**e**) pH 6.0 distal membrane ($M_D$) domain; (**f**) pH 9.2 $M_D$ domain. Local resolutions were estimated using cryoSPARC v3.3.2 and visualized using UCSF ChimeraX 1.8. FSC curves were generated using RELION-4.0 postprocessing.

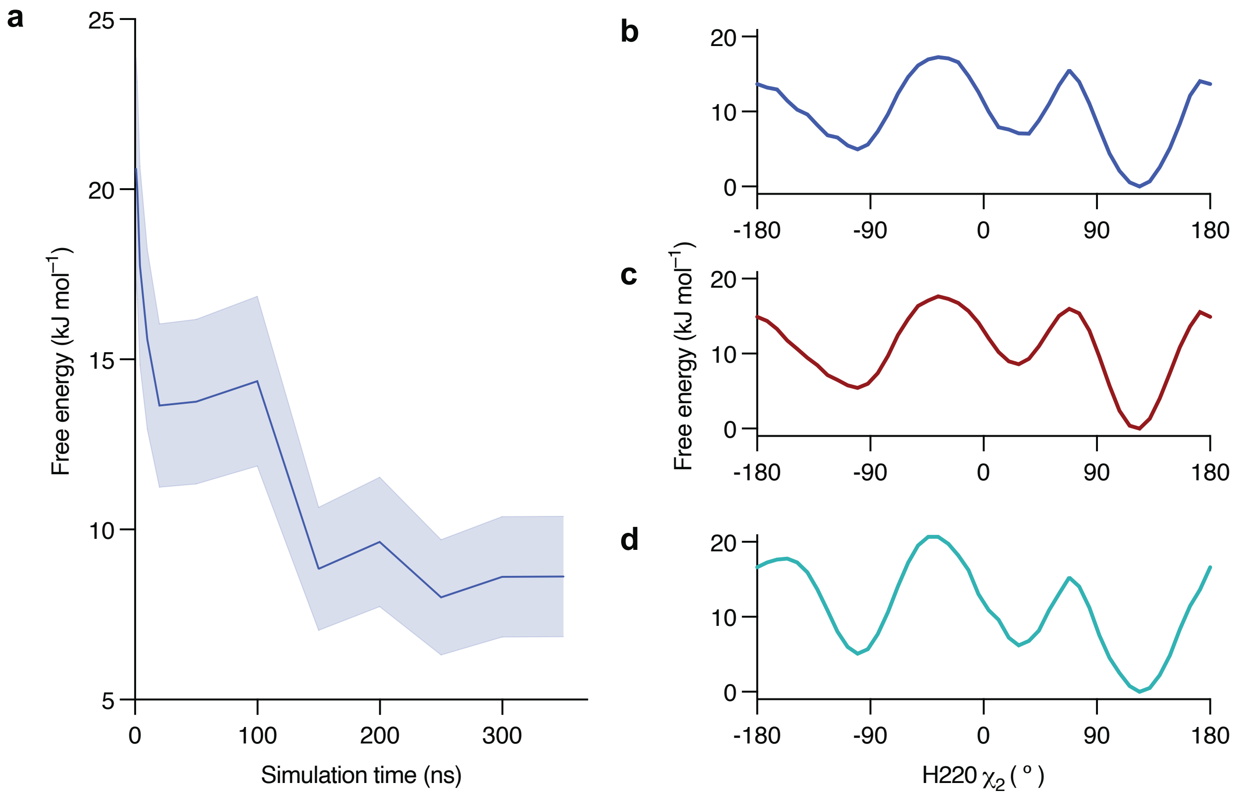


**Supplementary Fig. 7. Convergence of the computed free energy profiles for H220$^{ND4}$.** The analysis corresponds to the simulations carried out with H220$^{ND4}$ neutrally charged (protonated on N$\delta$, HSD) and the protonation state of other residues set to their predicted states at pH 6.0 (Fig. 4b). (**a**) Convergence of the free energy with simulation time, computed as the difference between $\chi_2$ of 123° and 42°. The statistical uncertainty from bootstrap analysis is shown as the shaded region. Variations in the free energy difference become smaller than the statistical error after ~150 ns, indicating convergence. (**b-d**) Free energy profiles obtained by splitting the full trajectory (380 ns) into three equally sized blocks. The similar overall shapes and relative free energies across these profiles and the profile for full trajectory shown in Fig. 4b (with standard deviations < 1.5 kJ mol$^{-1}$ in the region corresponding to experimental values 0° < H220 $\chi_2$ < 180°) support convergence and that the results are not strongly sensitive to initial configurations. Similar behaviour was observed for the other free energy profiles calculated.
